# Minimally invasive measurement of maternal transcripts enables predicting the developmental potential of mammalian zygotes

**DOI:** 10.64898/2026.06.02.726088

**Authors:** Akihiro Inoue, Nicole Cheung, Tadayuki Yamanouchi, Hideo Matsuda, Hajime Yoshioka, Hiroki Takeuchi, Mikiko Nishioka, Mari Yamamoto, Yubao Wei, Kazuya Houri, Hideo Sato, Renlong Guo, Asuka Kamio, Hisato Kobayashi, Tomohiro Kono, Kazuya Matsumoto, Kei Miyamoto

## Abstract

Maternal transcripts are stored in the oocyte cytoplasm during oogenesis and play a pivotal role in early embryonic development after fertilization. However, specific maternal transcripts that reflect the developmental potential of embryos have not been systematically identified, and the use of maternal transcript levels as an indicator of successful development has not been explored. Here, we link the maternal transcriptome to the zygote’s developmental potential by examining transcripts in a single polar body. The transcriptome of a zygote or an oocyte was highly similar to that of its accompanying polar body in mouse, cow, and human. We have identified a set of maternal transcripts whose expression levels fluctuate between poor- and good-quality zygotes. Specifically, *Sipa1*and *Zmym6* were identified as marker transcripts that accurately reflect the developmental potential of zygotes. Using these marker genes, combined with machine learning, the development of zygotes to the blastocyst stage was successfully predicted with more than 80% specificity as early as 12 hours after fertilization. Furthermore, our prediction platform significantly improved implantation rates and live births to term. Thus, we have demonstrated a minimally invasive method for identifying maternal transcripts associated with zygote developmental potential. Our developed prediction system provides a generalizable conceptual framework for human infertility treatment to reduce the risk of implantation failure by excluding embryos with low developmental potential, especially when early embryos are transferred, and for livestock propagation to assess selected expressed maternal trait-associated variants before embryo transfer.

## Introduction

Embryo selection remains a central challenge in reproductive biology, where the success rate of assisted reproductive technologies (ART) is profoundly affected by the quality of embryos: poor-quality embryos potentially lead to implantation failure and miscarriage. Current approaches largely rely on morphological assessment and, more recently, chromosomal screening techniques such as preimplantation genetic testing for aneuploidy (PGT-A). PGT-A is used to identify genetic abnormalities in embryos before transfer to the uterus, reducing the risk of miscarriage ^1^. While PGT-A assesses the genetic abnormalities of some embryonic cells, the detection of aneuploidy does not necessarily reflect the developmental potential of embryos ^2^. As a result, there is a growing need for minimally invasive strategies that can capture the underlying biological determinants of early embryogenesis. Many efforts are underway to judge the embryo’s viability. Morphokinetic assessment of preimplantation embryos using time-lapse imaging is clinically used with the development of artificial intelligence (AI)-based models ^3^. Metabolic profiling of spent culture media, namely metabolic fingerprinting, has been explored as a non-invasive biomarker of embryo fitness ^4^. However, the genetic and molecular biomarkers associated with embryonic developmental potential remain unclear.

Early embryonic development is predominantly governed by maternal factors deposited during oogenesis. Prior to zygotic genome activation (ZGA), which occurs at the 2-cell stage in mice and later stages in humans and cows, the embryo relies on maternally inherited transcripts and proteins to regulate key processes such as cell division and epigenetic reprogramming ^5^. Transcriptome studies revealed dynamic expression patterns of maternal transcripts during early embryonic development ^6^, and the spatiotemporal regulation of their translation is required for zygote development ^7^. Developmental failures of embryos are associated with abnormal expression of maternal transcripts ^8^. Functional experiments in which maternal genes were knocked out demonstrated their importance for development, leading to the identification of maternal-effect genes ^9–11^. In addition, recent advances in single-cell transcriptomics and developmental profiling have enabled increasingly precise characterization of embryonic cell states with RNA signatures ^12,13^. These reports suggest that expression of maternal transcripts may serve as a biomarker for predicting the developmental potential of embryos. However, due to the lack of appropriate experimental systems to unbiasedly measure maternal transcripts and simultaneously assess oocyte or zygote development, it remains unclear which maternal transcripts best predict the developmental potential of oocytes or zygotes.

In mammals, an oocyte arrests at prophase of meiosis I and, after resumption of meiosis, extrudes a first polar body (PB1). Then, a mature oocyte again arrests at metaphase of meiosis II (MII). Upon its fertilization with sperm, an MII oocyte extrudes a second polar body (PB2). Polar bodies (PBs) do not develop and are degraded when embryonic development proceeds. Therefore, it is possible to biopsy PBs without damaging their sibling oocytes or zygotes. By making use of these characteristics of PBs, PBs serve as a source of deducing genomic information of sibling oocytes, such as aneuploidy and SNPs ^14^, for clinical diagnostics ^15^. In addition, polar body genome transfer is used to prevent the transmission of mtDNA mutations that cause inherited mtDNA diseases ^16,17^. These studies use PB genomes, but PBs also share their cytoplasm with oocytes until cell division, suggesting that a PB may contain transcripts similar to those of oocytes. RNA-seq analysis of human PB1 suggests that PB1 reflects the oocyte transcript profile ^18^. Reverse transcription-polymerase chain reaction (RT-PCR) analyses revealed that at least highly abundant maternal transcripts are detected in PB2 ^19^. When aiming to develop a method to accurately and reproducibly detect maternal transcripts in PBs, PB2 might be a better choice than PB1. This is because ovulated MII oocytes contain those that passed hours after PB extrusion, during which transcripts in PB1 may be degraded. Indeed, ovulated mouse MII oocytes sometimes lack intact, visible PB1, suggesting that PB1 may disintegrate within hours. In contrast, PB2 extrusion can be closely monitored in embryos after *in vitro* fertilization (IVF). RNA-seq analysis of pooled PB2s suggested that almost all transcripts in PB2s were expressed in zygotes ^20^. However, it is unclear whether the transcriptome of a zygote is mirrored in its accompanying single PB2. Moreover, the relationship between the transcriptional pattern in PB2 and the developmental potential of its sibling zygote has not been studied.

Here, we establish a minimally invasive transcriptomic strategy for inferring embryonic developmental potential from PB2-derived maternal RNA signatures. Our single-cell RNA-seq (scRNA-seq) data revealed the mouse PB2 transcriptome and showed that PB2 mirrored the transcription profile of its sibling zygote. We then investigated the relationship between the PB2 transcriptome and the developmental potential of its sibling zygote, leading to the identification of transcripts enriched in PB2 extruded from zygotes that develop into blastocysts, or from those that arrest before the blastocyst stage. Among these transcripts, we selected a few genes whose expression in PB2 is used to predict development. Using our method, the development of mouse zygotes to blastocysts can be predicted with 80.8% specificity. Furthermore, machine learning-based prediction from maternal transcript measurements enabled rapid assessment of zygote developmental potential within a day (as early as 12 hours after fertilization), resulting in significant improvement in development to term. Thus, our study presents a conceptual framework with predictive validation in mice for future application in human ART and livestock propagation.

## Results

### Maternal transcripts enriched in a zygote are expressed in a second polar body

To profile transcripts in a polar body, we examined transcriptomes of a PB2 and its sibling zygote by single cell RNA-seq using ERCC spike-in references for normalization ^21,22^. This approach resulted in high read coverage and alignment rates to the mouse genome (Extended Data Table S1). Principal component analysis (PCA) using all detected transcripts showed a separation between PB2s and zygotes (Extended Data Fig. 1a), reflecting the difference in the number of detected transcripts due to the large volume of a zygote. Indeed, zygotes showed 13,211 ± 313 transcripts while 9,096 ± 615 transcripts were detected in PB2s (Fig. 1a, mean ± SD). Then, we compared the type of transcripts between PB2s and zygotes. In all replicates, more than 96% of PB2 transcripts were also detected in zygotes (Fig. 1a). The transcripts detected both in PB2s and zygotes (shared transcripts in Fig. 1a) represented abundantly expressed transcripts in zygotes, while transcripts detected only in zygotes, but not in PB2s, were lowly expressed (Fig. 1b). These results suggest that abundantly expressed maternal transcripts are preferentially detected in PB2s. The transcriptomes of the shared maternal transcripts were highly similar among PB2s and zygotes (Fig. 1c), and expression levels of the shared transcripts in PB2s were well correlated with those in zygotes (Fig. 1d, R=0.92). Gene ontology (GO) analysis suggested that the shared transcripts were categorized into DNA repair and RNA processing (Fig. 1e), which resembled GO terms of mouse MII oocytes ^23^. On the other hand, transcripts uniquely expressed in zygotes did not show a clear enrichment of specific GO terms (Extended Data Fig. 1b). Furthermore, known maternal effect genes ^24^ were identified as shared transcripts and equally expressed both in zygotes and PB2s (Extended Data Fig. 1c). Together, abundantly expressed maternal transcripts are often detected in PB2s, whose expression levels reflect those in zygotes.

**Fig. 1.**
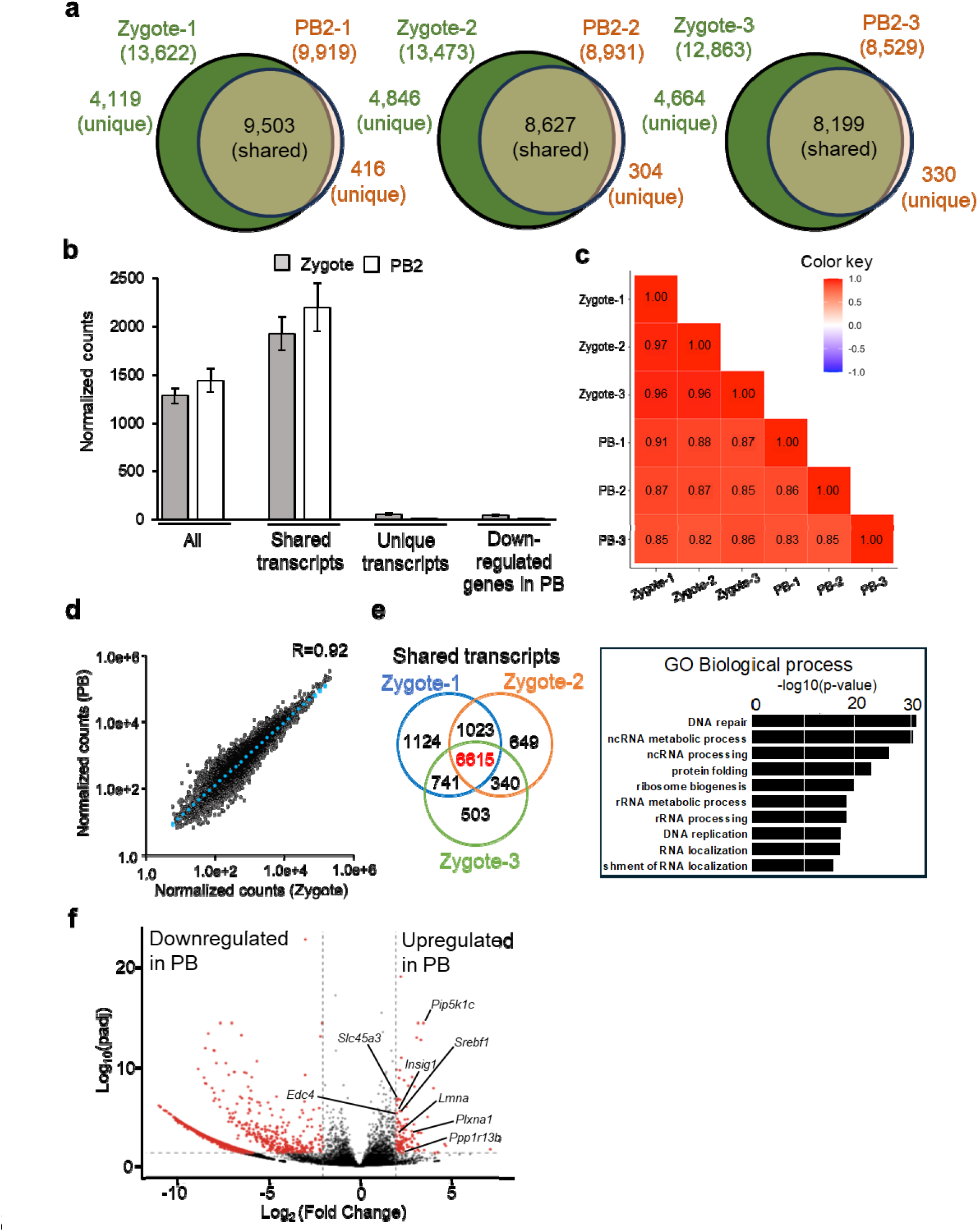
Polar body transcriptomes reflect maternal transcripts in sibling zygotes in mice. **a**, Venn diagrams showing the overlap of detected transcripts between PBs and their sibling zygotes. A substantial proportion of transcripts were shared across all samples (N = 3 biological replicates). Transcripts with counts >5 were regarded as expressed genes. Unique transcripts are those that were present in only one of the PB and Zygote samples. **b**, Averaged normalized counts of all transcripts, shared and unique transcripts, and downregulated transcripts in PB2s (**f**). Mean ± SD. N = 3 biological replicates. **c**, Correlation coefficients of transcriptomic profiles among zygotes and PB2s. **d**, A scatter plot showing the correlation of expression levels of shared transcripts between zygotes and PB2s and zygotes. **e**, A Venn diagram showing the overlap of shared transcripts (**a**), and 6,615 transcripts were detected as commonly expressed in all samples. GO terms for those transcripts are shown on the right. **f**, A volcano plot showing 125 upregulated transcripts (FC≧2, padj<0.05), and 1,809 downregulated genes (FC≦-2, padj<0.05). Genes that were found as maternal mRNAs degraded in zygotes ^25^ are indicated.

As shown, PB2s can be used to detect the maternally enriched transcripts in zygotes. However, it is still possible that a certain set of transcripts are enriched in zygotes or PB2s. We then identified differentially expressed genes (DEGs) by normalizing the difference of RNA abundance between zygotes and PB2s using the spike-in RNA (Fig. 1f); 125 upregulated and 1,809 downregulated genes in PB2s were found (padj<0.05, log2FoldChange≧2). The downregulated genes in PB2 showed much lower expression levels than the overall average (Fig. 1b), suggesting that their lower abundance in maternal cells hampered reproducible detection in PB2. The upregulated transcripts in PB2s might be related to the active uptake of maternal transcripts into PB2s, presumably due to the asymmetric distribution and/or for their degradation. To test the idea of active uptake of specific maternal RNAs for degradation, we first examined whether maternally degraded transcripts were enriched in the upregulated transcripts in PB2s. Still, only a few genes were overlapped between PB2-upregualted and maternally degraded genes (Extended Data Fig. 1d) ^25^. Next, we experimentally blocked the release of PB2s using Cytochalasin B (Extended Data Fig. 1e). However, the degradation of maternal transcripts, whose expression greatly drops by the 2-cell stage ^25^, was not affected, as judged by RT-qPCR analyses of 2-cell embryos (Extended Data Fig. 1f). Therefore, we did not obtain evidence that PB release is relevant to the degradation of maternal transcripts. In conclusion, expression levels of abundantly accumulated maternal transcripts in zygotes can be assessed by examining their levels in PB2s.

### Identification of maternal transcripts associated with the developmental potentials of mouse zygotes

We sought to identify maternal transcripts associated with the high- or low-developmental potential of zygotes. Because the expression pattern of maternal transcripts in zygotes was well reflected in PB2s (Fig. 1), we designed the experimental system in which a PB2 was collected from a zygote in a minimally invasive way and the manipulated zygote was further incubated to monitor the development (Fig. 2a). The stringent criteria were applied for collecting uniform PB2s (see relevant methods). First, to minimize experimental variation and individual differences, we tested three mouse strains. Among the strains tested, DBA/2 achieved high fertilization efficiency, and almost half of the *in vitro* fertilized embryos developed to the blastocyst stage (Extended Data Fig. 2a), thereby maximizing the efficiency of good and bad PB sampling. Culturing embryos in individual spots did not affect development to the blastocyst stage, as compared to culturing a group of embryos in individual spot (Extended Data Fig. 2b). Furthermore, the PB biopsy during 7-9 hpi did not affect development to the blastocyst stage (Extended Data Fig. 2c). Using these optimized conditions, PB2s collected from embryos that developed to blastocysts were classified as good PBs, while those from embryos that were arrested at the morula stage before reaching blastocysts were as bad PBs (Extended Data Fig. 2d). Then, gene expression between good and bad PBs was compared by single cell RNA-seq. The number of detected transcripts was similar between good and bad PBs (Fig. 2b). Transcripts of known maternal effect genes were reproducibly detected both in good and bad PBs (Fig. 2c). At a global level, good and bad PBs showed very similar transcriptomes, and the transcriptomes of good PBs were not separated from those of bad PBs by PCA (Fig. 2d,e). These results suggest that good and bad PBs cannot be distinguished at a global expression level in the mouse. We then identified DEGs between good and bad PBs (113 upregulated genes in good PBs and 80 downregulated genes in good PBs; padj<0.05, log2FoldChange≧1) (Fig. 2f). Genes related to metal iron-binding were enriched in DEGs, as revealed by GO analysis (Extended Data Fig. 2e). Ingenuity Pathway Analysis (IPA) also suggested the disturbance of iron homeostasis (Extended Data Fig. 2f). Among the identified DEGs, 23 genes were also shown as those related to the developmental potential of human oocytes ^26^ (Fig. 2g, h). These genes can serve as promising biomarkers for predicting the developmental potential of mammalian oocytes/zygotes. Taken together, although embryos that developed normally to the blastocyst stage did not show apparently different transcriptomes at the zygote stage from those arrested before the blastocyst stage in our experimental setup, a set of specific genes were differentially expressed.

**Fig. 2.**
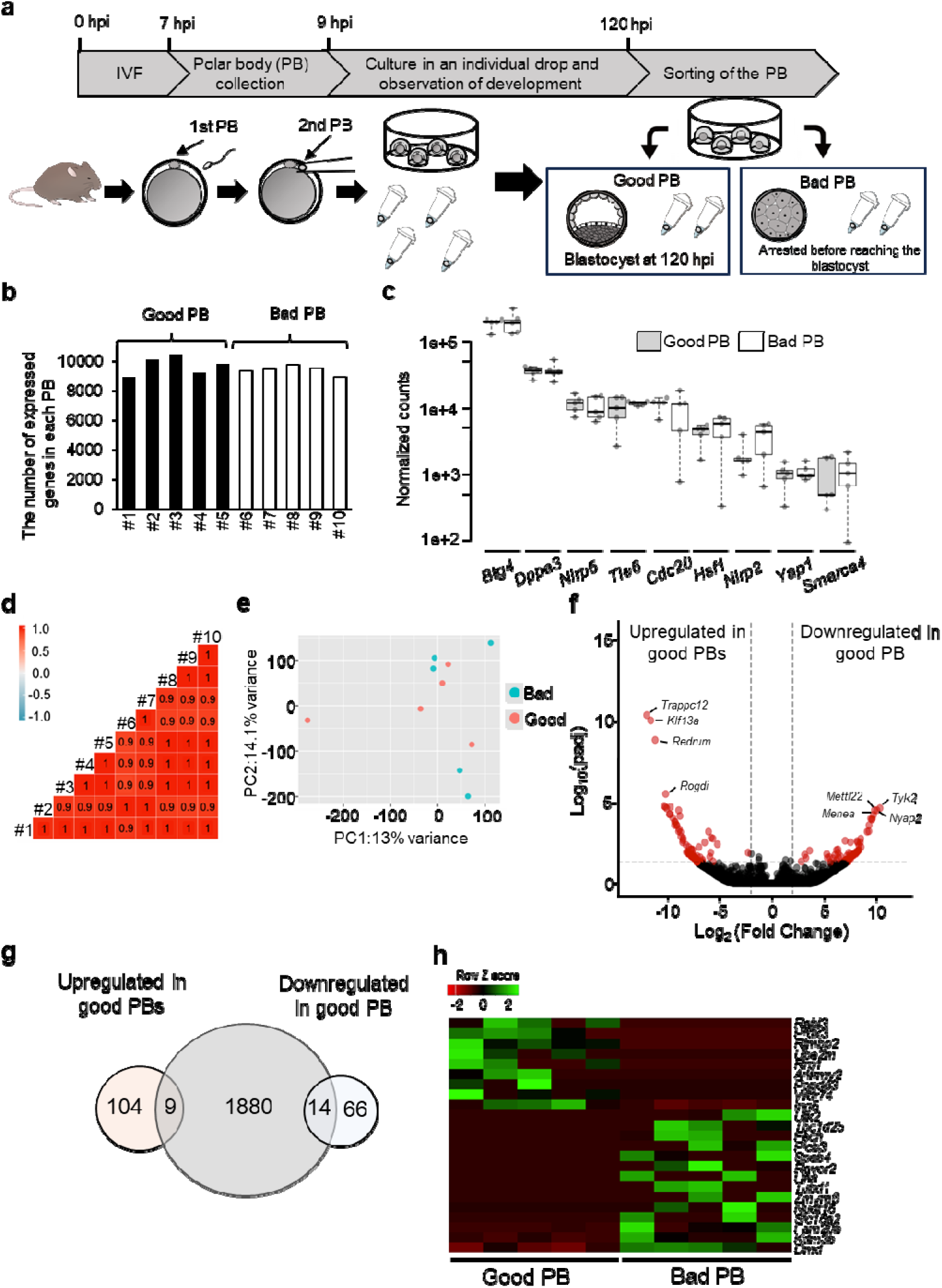
Identification of candidate biomarker maternal transcripts in polar bodies, whose expression is associated with the embryonic developmental potential. **a**, Experimental design for PB2 sampling and sorting Good- vs Bad-PBs. PB2s were biopsied from zygotes during 7-9 hpi, followed by *in vitro* culture in an individual drop. After 5 days of culture, PB2s associated with embryos that normally developed to the blastocyst stage were classified as Good PB. In contrast, PB2s with embryos that were arrested before the blastocyst stage as Bad PB (Extended Fig. 2d). **b**, The number of detected transcripts in Good and Bad PBs (counts >5). **c**, Normalized counts of representative maternal genes in Good and Bad PBs. N = 5 (Good PB) and 5 (Bad PB) biological replicates. The expression levels did not differ between good and bad PBs, as determined by T-test (P > 0.05). **d**, Correlation of transcriptomes among Good and Bad PBs. **e**, PCA of the PB2 transcriptomes. **f**, A volcano plot showing 80 upregulated and 113 downregulated genes in Good PBs (|log2FC|≧1, padj<0.05). **g**, A Venn diagram showing the overlap of up- and down-regulated genes in Good PBs with oocyte-expressed genes whose expression fluctuates depending on developmental potentials in humans ^26^. **h**, Heatmap of selected 23 DEGs identified in (**g**) as overlapping with human developmentally related genes.

### Maternal transcripts stored in bovine oocytes

We next aimed to identify maternal transcripts associated with the developmental potential of bovine zygotes. Since *in vitro* maturation (IVM) is widely used in bovine reproduction, we first examined its effects on zygote and PB2 transcriptomes collected from the same cows using scRNA-seq (Fig. 3a). Bovine zygotes contained 12,109 ± 406 transcripts, whereas the accompanying PB2s showed 5,258 ± 1,444 transcripts (Fig. 3b, Extended Data Table S1, mean ± SD). The hierarchical clustering and PCA showed clear separation between zygotes and PB2s, although *in vivo* and *in vitro* zygotes were not clearly clustered (Extended Data Fig. 3a,b). Furthermore, few DEGs were detected between *in vitro* and *in vivo* zygotes (Extended Data Fig. 3c; 0 DEGs with padj<0.05, log2FoldChange≧2: 264 DEGs with p<0.05, log2FoldChange≧2). GO analysis revealed enrichment of cell adhesion-related terms (Extended Data Fig. 3c). These results suggest that *in vitro* maturation does not profoundly affect mRNA expression profiles, although fluctuations in transcriptomes may occur in some samples.

**Fig. 3.**
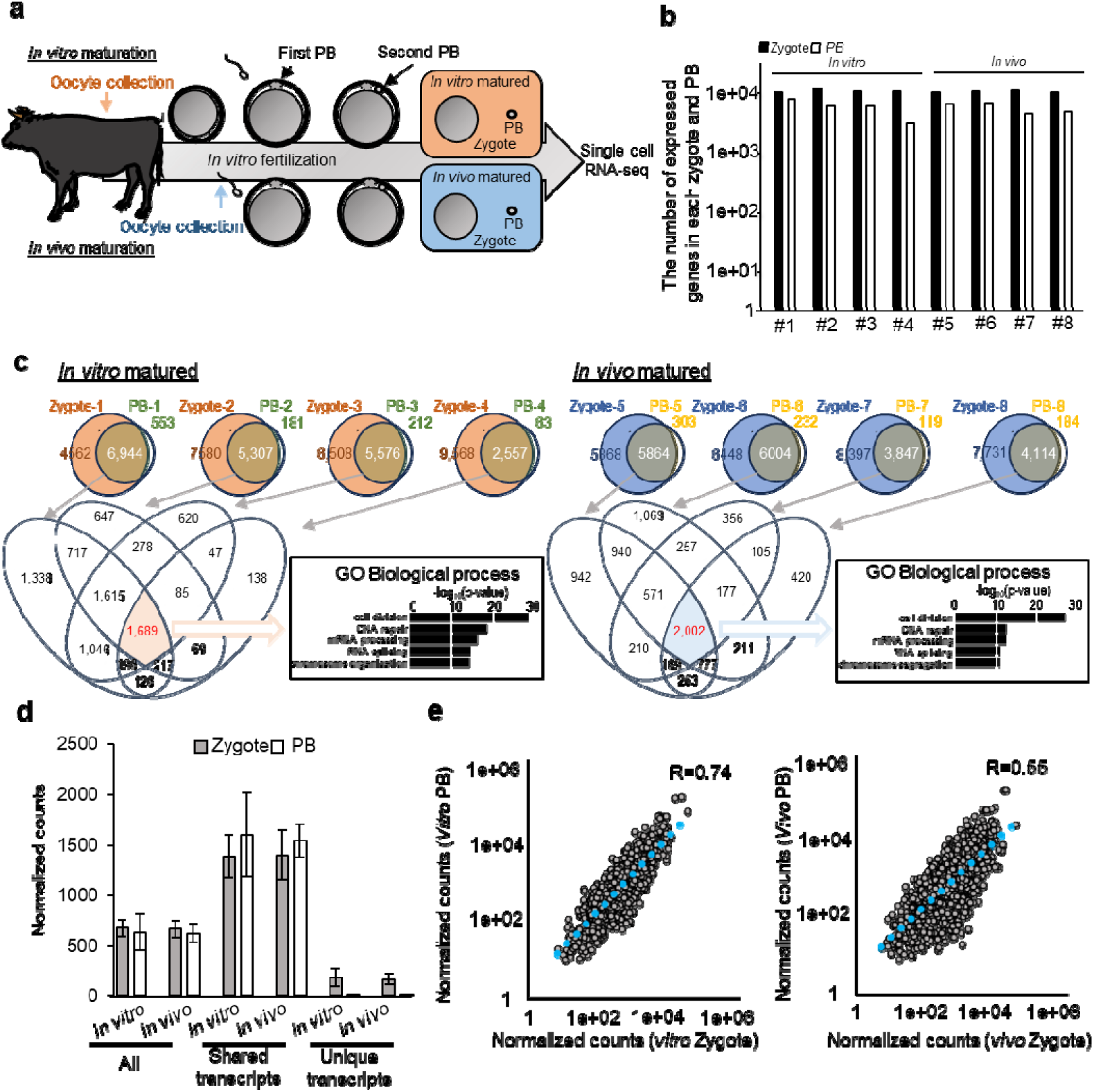
Transcriptomes of polar bodies reflect their associated zygotes derived from *in vitro* and *in vivo* matured bovine oocytes. **a**, Experimental design of bovine zygote and PB2 sampling from *in vitro*– and *in vivo*–matured oocytes using the same cows. **b**, The number of detected transcripts in zygotes and PBs (counts >5). **c**, Venn diagrams showing the overlap of detected transcripts between *in vitro*- and *in vivo*-matured zygotes and PBs, with associated GO biological processes. **d**, Normalized counts of all samples, shared and unique transcripts. Mean ± SD. N = 4 biological replicates. **e**. Correlation of transcript expression levels between *in vitro*- and *in vivo*-matured zygotes and PBs.

We then compared transcriptomes between PB2s and their sibling zygotes. Strikingly, more than 95% of transcripts detected in PB2s were also present in their sibling zygotes (Fig. 3c). Commonly detected transcripts (1,689 for *in vitro* maturation and 2,002 for *in vivo* maturation) were enriched with the same GO terms (Fig. 3c), indicating a substantial overlap in transcriptional composition. This high degree of overlap suggests that PB2s retain a representative subset of maternal transcripts present in zygotes. These shared transcripts were highly expressed in zygotes, whereas transcripts detected only in zygotes were lowly expressed (Fig. 3d, unique transcripts), indicating that the highly expressed maternal transcripts are readily detected in PB2s. Expression profiles of shared maternal transcripts showed strong correlations between PB2s and zygotes (Fig. 3e). Taken together, these results indicate that PB2 transcriptomes can serve as a minimally invasive proxy for assessing the molecular state of zygotes in both *in vitro*- and *in vivo*-matured oocytes.

### Identification of maternal transcripts associated with the developmental potential and genetic variability in bovine zygotes

We next investigated maternal transcripts associated with the developmental potential in cows by taking advantage of the PB-based method developed in mice, shown in Fig. 2. PB2s were isolated from bovine zygotes and classified into three groups based on IETS morphological criteria ^27^ (Fig. 4a, see Methods for more details). Neither hierarchical clustering nor PCA revealed clear separation among groups, but the transcriptomes of very good PB clustered relatively closely (Fig. 4b,c, circle), indicative of similar transcriptomic profiles in the high-quality embryos. Expression levels of known maternal effect genes showed no substantial differences among PB2s derived from different morphological grades (Fig. 4d). These observations correspond to the results of mouse zygotes (Fig. 2). Then, we examined DEGs between Bad PB and Very Good PB groups, leading to the identification of 73 upregulated and 658 downregulated genes in the Very Good PB group (padj < 0.05, |log2FC| ≥ 2) (Fig. 4e). GO and IPA revealed that Bad PBs were characterized by activation of meiotic dysregulation, DNA damage, and stress response pathways, whereas Very Good PBs exhibited enrichment of pathways related to metabolic homeostasis and protein quality control (Extended Data Fig. 4a,b). Notably, genes associated with oocyte meiosis were expressed at lower levels in Very Good PBs, whereas they were upregulated in Bad PBs and Moderately Good PBs (Fig. 4f). This suggests that elevated oocyte meiosis signatures in PB2 reflect a transcriptional residual of meiotic dysregulation in zygotes, potentially associated with defective chromosome segregation and/or embryonic cell cycle regulation. Interestingly, cross-species comparison between mouse and bovine PB transcriptomes revealed that only a limited number of differentially expressed genes were conserved. However, five genes showed consistent expression patterns across species in both Good and Bad PB groups (Extended Data Fig. 4c, *Commd4*; *Tmem120a*; *Aacs*; *Zmym6*; *Mfsd11*). These genes may represent evolutionarily conserved regulators associated with embryonic development. Furthermore, scRNA-seq of PB2 enabled SNP detection from sequencing reads. Although detectable SNPs were limited to loci covered by RNA-seq reads, several known variants associated with economically important traits, such as carcass weight and fertility, were successfully identified ^28–30^ (Fig. 4g). Notably, the NCAPG variant (c.1326 T>G), associated with carcass weight ^28^, showed high concordance between PB2s and corresponding zygotes (Extended Data Fig. 4d). This highlights the usage of PB analysis for simultaneous assessment of both developmental competence and genetic traits. Importantly, such PB-biopsied and PB-sequenced embryos were transferred, and healthy calves were born (Fig. 4h). Furthermore, the detected SNP at the *NCAPG* locus was maintained in the resulting calves as G/T (Extended Data Fig. 4e), supporting feasibility at maternally expressed, trait-associated loci. Collectively, PB-based transcriptomic analysis provides a minimally invasive approach to evaluating bovine developmental competence and demonstrates the feasibility of detecting selected trait-associated SNPs at maternally expressed loci.

**Fig. 4.**
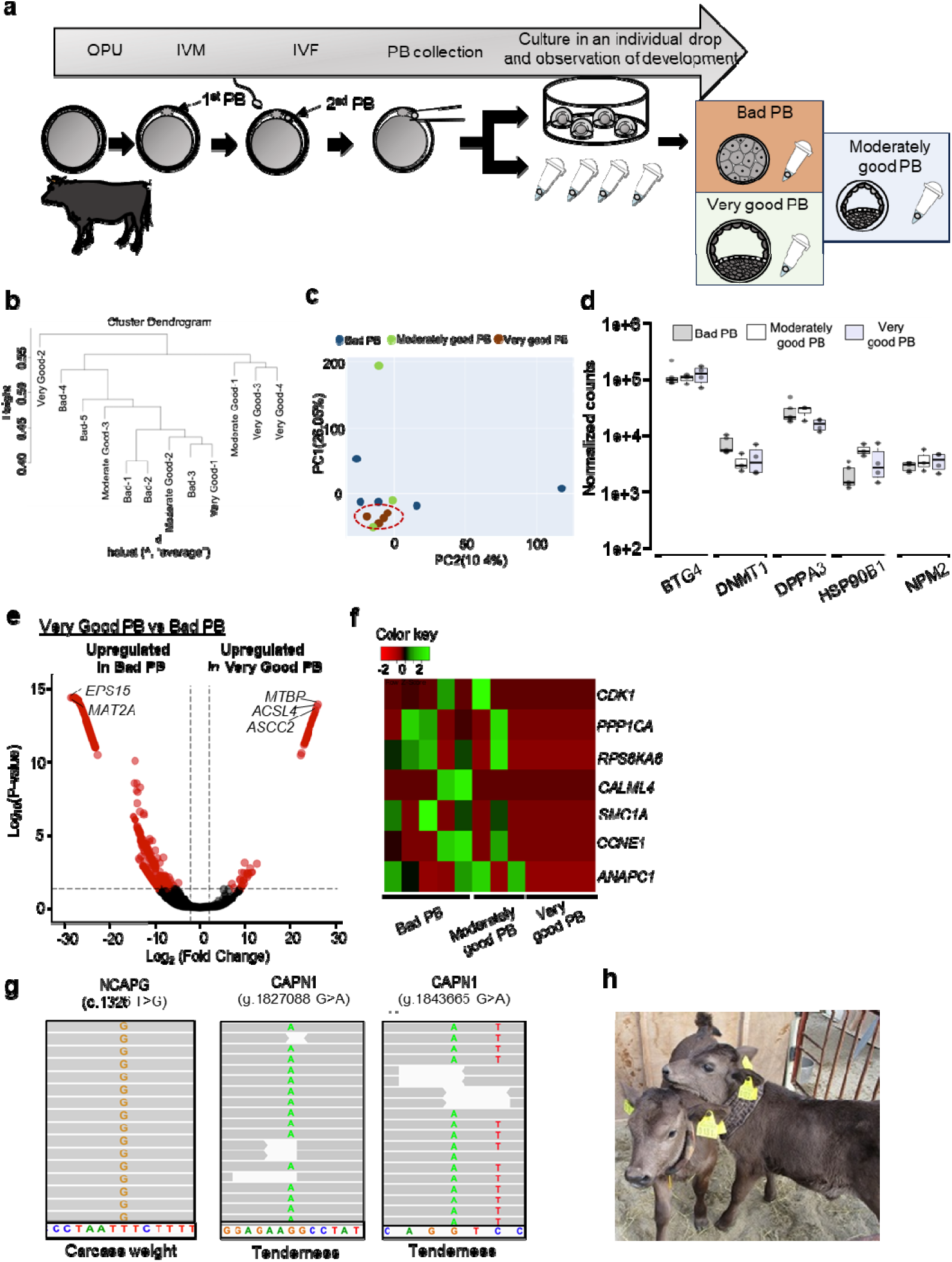
Polar body transcriptomes distinguish embryo quality states and reveal candidate markers of developmental competence in the cow. **a**, Experimental workflow showing OPU, IVM, IVF, PB collection, and subsequent embryo culture with developmental assessment. Very good PB: PBs collected from embryos that developed to blastocysts (code 1 quality according to the IETS standard). Moderately good PB: PBs collected from embryos that developed to blastocysts (code 2), in which abnormalities in the outline, size and color of individual cells were observed. Bad PB: PBs from embryos arrested before the blastocyst stage. **b**, Hierarchical clustering of PB transcriptomes. **c**, A PCA plot illustrating the separation of PB samples across the three groups. Transcriptomes of very good PBs were similar (indicated by a circle). **d**, Boxplots showing normalized expression levels of representative maternal effect genes. N = 4 (very good PB), 3 (moderately good PB), and 5 (bad PB). The expression levels did not differ among samples, as determined by ANOVA (P > 0.05). **e**, A volcano plot identifying DEGs associated with developmental competence (very good PBs vs Bad PBs) (|FC|≧2, padj<0.05); gene names of the most fold-changed genes were highlighted. **f**, Heatmap of selected genes involved in oocyte meiosis and early embryonic development, showing group-specific expression patterns. **g**, Detection of genetic variants in PB-derived RNA-seq, which are related to meat quality and conception rate. **h**, The born calf after PB biopsy and examination by RNA-seq.

### Maternal transcripts enriched in an unfertilized oocyte are detected in a human PB1

We next asked if a variety of maternal transcripts are detected in a human polar body. As shown above, a PB2 was used as a source for quantifying maternal transcripts in mouse and bovine zygotes, but it is ethically difficult to collect PB2s from normal human zygotes, most of which are used for infertility treatment. Therefore, we sampled PB1s from unfertilized surplus MII oocytes to assess maternal transcript expression. Human oocytes were retrieved by follicular puncture from 6 patients and *in vitro* matured to the MII stage ^31^. Oocytes and their sibling PB1s were each analyzed by single cell RNA-seq (Extended Data Fig. 5a); MII oocytes *in vitro* matured on the day of oocyte collection were described as MII (MII-1-3) and PB (PB-1-3), and those *in vitro* matured to the MII stage till the next day of oocyte collection as delayed MII and PB (dMII-1-3 and dPB-1-3) ^32^. At the transcriptomic level, oocyte and polar body samples were clustered separately (Extended Data Fig. 5b, c), reflecting the number of detected transcripts (Fig. 5a). The expression pattern of oocytes highly resembled each other, while that of polar bodies was less well correlated, although still exhibiting the reasonable correlation (Extended Data Fig. 5d). On average, an oocyte and polar body contained 16,295 ± 419 and 10,768 ± 1,432 transcripts (normalized gene counts more than 5), respectively and 93.2 ± 1.0% of the polar body transcripts were also expressed in an oocyte (n = 6, mean ± SD), suggesting that a myriad of maternal transcripts are stored in a PB1. When PB-1-3 were compared to dPB-1-3, the number of detected transcripts were not much different (Fig. 5a; 10,683 ± 1,985 and 10,854 ± 385, respectively). We next identified 5,880 transcripts that were commonly expressed in all PB and MII, and 6,812 transcripts were in all dPB and dMII (Fig. 5a). GO analysis and IPA identified the same set of biological processes, highlighting a high level of similarity in biological activities shared between the two groups (Fig. 5a and Extended Data Fig. 5e). These shared genes between oocytes and polar bodies represented abundantly accumulated maternal transcripts as compared to genes that were uniquely detected in MII oocytes or PBs (Fig. 5b). We then examined expression levels of known maternal effect genes in humans ^24^. All maternal effect genes tested were detected in both PB-1-3 and dPB-1-3, although the expression level of *TPIP13* was variable (Fig. 5c). To note, *TRIP13* showed the weakest expression in oocytes among the examined genes, implying that expression levels of maternal transcripts in oocytes may influence detection efficiency in PBs. We next identified DEGs between MII (MII-1-3 and dMII-1-3) and PB (PB-1-3 and dPB-1-3) by normalizing with ERCC spike-in reference (Fig. 5d). In the comparison between MII versus PB, not many DEGs were identified (Fig. 5d, 104 upregulated and 218 downregulated genes in PB [padj < 0.05, |log2FC| ≥ 2]). In contrast, 4,975 and 807 genes were upregulated in dPB and dMII, respectively, in the delayed condition (Fig. 5d; padj < 0.05, |log2FC| ≥ 2). Furthermore, highly correlated expression levels of shared transcripts were found in MII versus PB when sampled promptly, while the degree of correlation decreased between dMII and dPB (Fig. 5e). These results suggest that the prolonged culture of oocytes alters transcriptomes in oocytes and/or PBs, diminishing the mirroring characteristics of the PB to assess maternal transcripts. Among the PB-upregulated genes, highly abundant maternal transcripts, such as *TUBB8*, *KHDC3L* and *TLE6*, were found (Fig. 5d). Therefore, in general, maternal transcripts in PB1s reflected their expression levels in MII oocytes, but the prolonged culture of oocytes resulted in enrichment of some abundant maternal transcripts in polar bodies. Abundant *TUBB8* transcripts enabled the detection of known mutation sites that are associated with infertility (Fig. 5f) ^33^, indicating that, in theory, PB transcripts can also be used for genetic diagnosis, such as preimplantation genetic testing for monogenic disorders. In conclusion, abundant maternal transcripts are reproducibly detected in the human PB1 when collected immediately during meiosis.

**Fig. 5.**
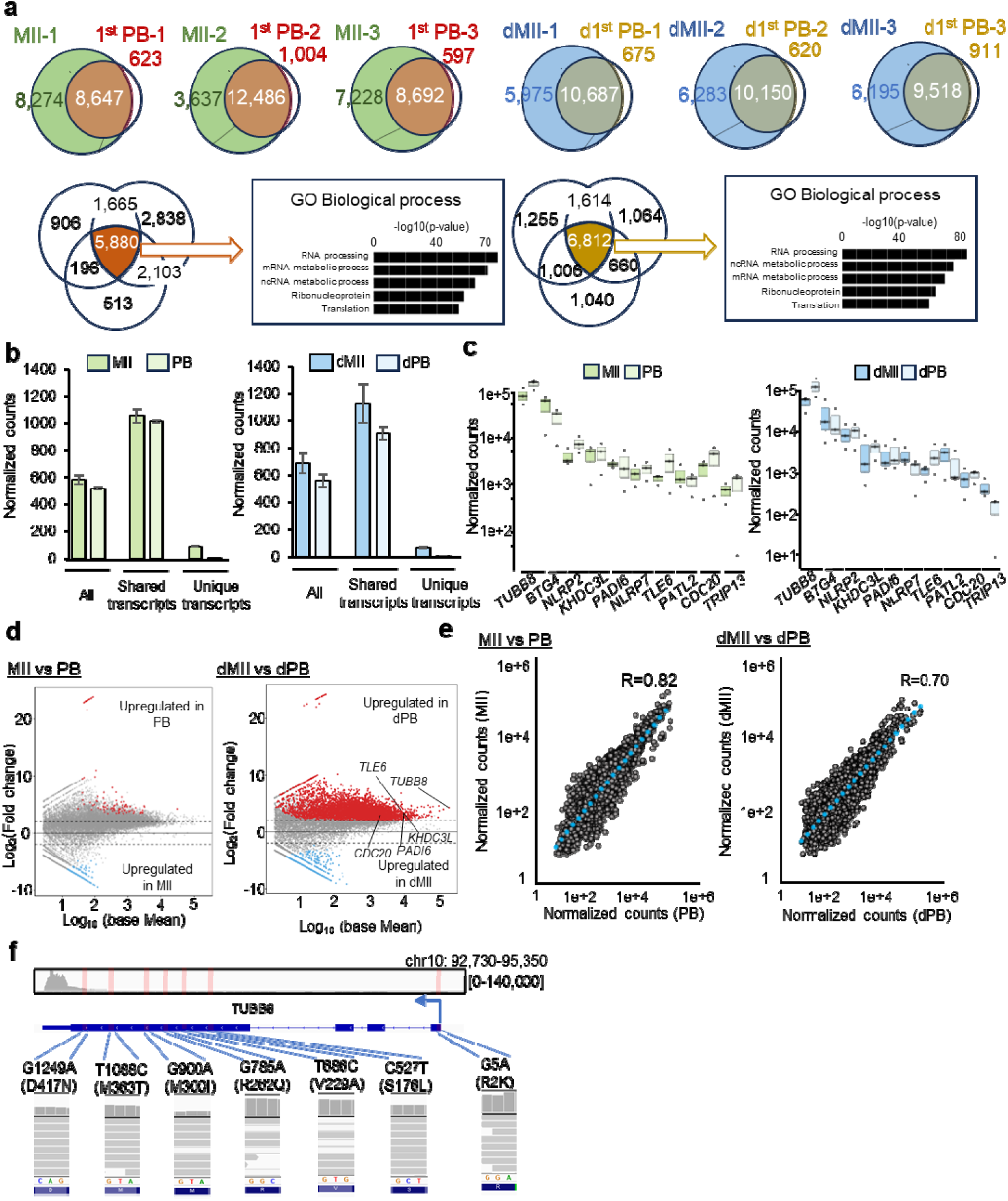
Transcriptomes of 1^st^ polar bodies reflect their associated MII oocytes in the human. **a**, Venn diagrams showing overlap of detected transcripts between individual MII oocytes and their corresponding first polar bodies (PB1). Numbers indicate shared and unique transcripts in each pair. dMII and dPB mean oocytes delayed in their maturation. GO enrichment analysis of shared transcripts, indicating enrichment for RNA processing, mRNA metabolic process and translation-related pathways. N = 3 (MII and PB) and 3 (dMII and dPB). **b**, Expression levels of all, shared, and unique transcripts in MII and dMII conditions. Mean ± SD. N = 3 biological replicates in each group. **c**, Boxplots showing normalized expression levels of representative maternal effect genes. N = 3 biological replicates in each group. The expression levels did not differ between MII oocytes and PBs, as determined by the T-test (P >0.05). **d**, MA plots showing DEGs between MII oocytes and PBs and those between dMII and dPB (|FC|≧2, padj<0.05), with significantly upregulated genes in PBs highlighted in red and upregulated in MII oocytes highlighted in blue. Some maternal effect genes identified as DEGs are indicated. **e**, Correlation of transcript expression levels between MII and PB and those between dMII and dPB. **f**, Representative sequencing reads showing detection of single-nucleotide variants of *TUBB8* in PB RNA-seq data.

### Maternal transcript measurement using a polar body enables rapid and accurate prediction of the embryonic developmental potential

Our results in mice, cows, and humans demonstrate that PB can serve as a minimally invasive indicator of maternal transcript expression, which can be used to identify biomarker transcripts associated with developmental potentials of zygotes (Fig. 2 and 4). We therefore aimed to identify predictive markers associated with successful embryonic development and to develop a transcript biomarker-based approach to predict developmental outcome. To this end, predictive markers were selected through a 3-step screening strategy designed to minimize technical variation and inter-individual differences while ensuring reproducibility (Fig. 6a). In screening step 1, 29 candidate genes were selected from DEGs identified by scRNA-seq analysis as performed in Fig. 2 (Fig. 6b). The selection of those genes was based on the criteria of reported functions in early embryonic development ^34,35^, identification across species (Fig. 2g), or abundant and reproducible expression in our RNA-seq dataset (Extended Data Fig. 6). In screening step 2, these candidates were first tested by RT-qPCR using a single PB with a few number of biological replicates for the initial screening (Extended Data Fig. 7a). When the detection rate was less than 20% by single PB RT-qPCR, those transcripts were not further analysed (Extended Data Fig. 7a). The RT-qPCR confirmation process was further repeated using cDNA-amplified PB samples (Extended Data Fig. 7b, c). Genes showing the strong association with embryonic developmental outcomes were selected, and further validated by single PB RT-qPCR (Fig. 6c, Extended Data Fig. 7d, e), leading to the identification of *Cdk2ap1*, *Sipa1*, and *Zmym6* as candidate markers associated with developmental competence (Fig. 6c). In addition, *Dppa3*, one of the most abundantly accumulated maternal transcripts whose expression does not differ between good and bad PBs, was included to account for potential differences in transcript abundance due to the PB2 size, resulting in a final set of four predictive markers.

**Fig. 6.**
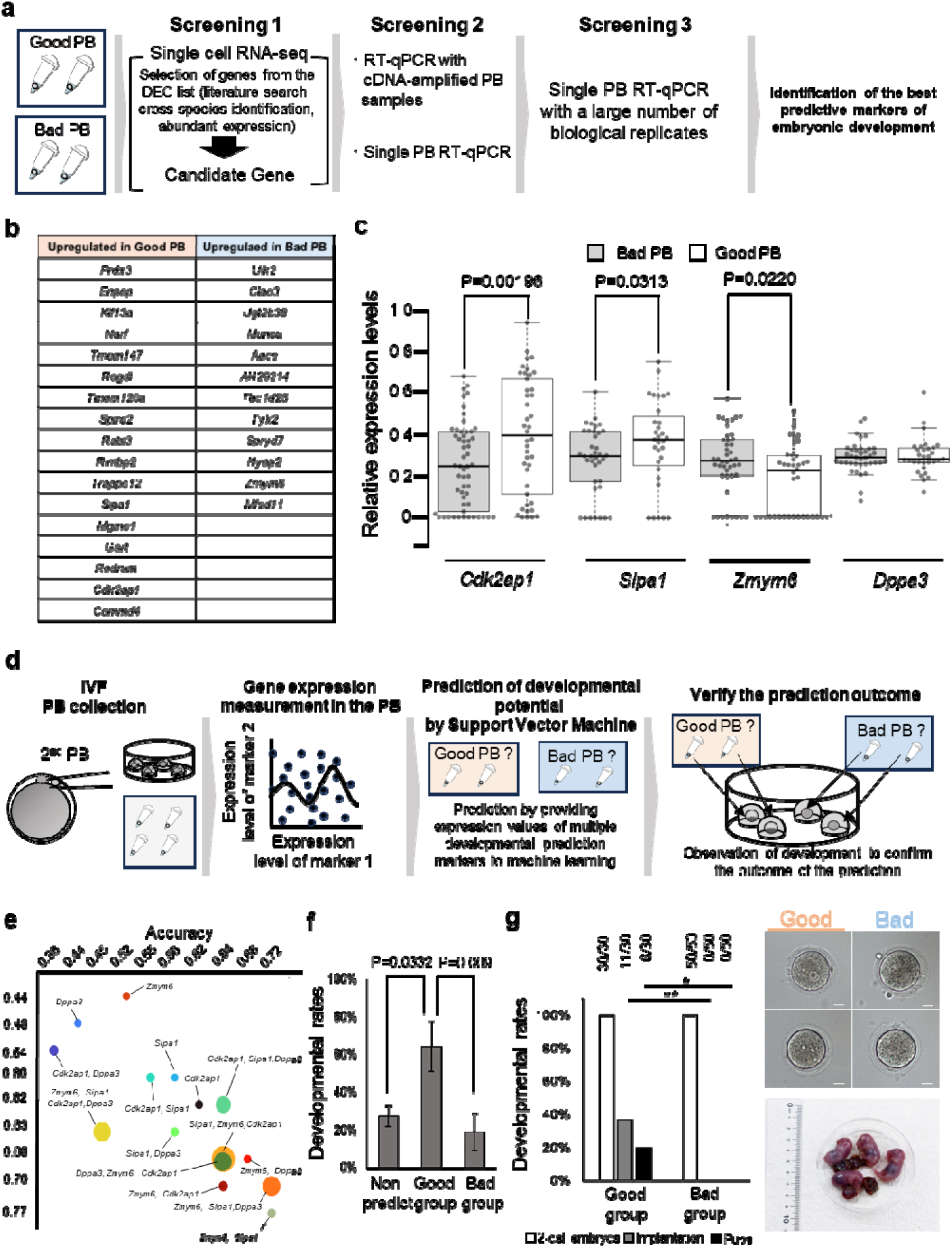
A predictive model using polar body transcripts enables classification of embryonic developmental potentials. **a**, Workflow for identifying biomarker transcripts. Good and Bad PB were collected by following the scheme shown in Fig. 2. Using the DEG list in Fig. 2f, screening for the best predictive biomarkers was performed. **b**, Upregulated gene lists in Good- and Bad- PB groups after the screening 1 (**a**). **c**, Box plots showing the relative expression levels of final candidate biomarkers as judged by single PB RT-qPCR: *Cdk2ap1, Sipa1, Zmym6, and Dppa3*. T-test was performed. **d**, Scheme for establishing machine learning-based developmental prediction model, including PB collection, gene expression measurement of *Cdk2ap1, Sipa1, Zmym6, and Dppa3*, PB classification, and verification. **e**, Various combinations of marker transcript expression were tested to establish a prediction model. X axis: accuracy, Y axis: Area Under the Curve (AUC), Circle size: probability of misclassification. **f**, Prospective cohort study to assess the embryonic development to the blastocyst stage. Percentages of development to the blastocyst stage in each group are shown. Developmental prediction and classification were performed at the zygote stage. Mean ± SE. N = 4 experimental replicates. Statistical significance was evaluated by the Chi-square test. **g**, Prospective cohort study to assess the embryonic development to term. Percentages of development to pups and implantation rates are shown in each group. Developmental prediction and classification were performed at the zygote stage, and the classified 2-cell embryos were subjected to embryo transfer. Representative images of the Good and Bad group embryos are shown. Furthermore, the newborn mouse pups obtained from the Good group are shown. Scale bar, 20 μm. N = 3 biological replicates. Fisher’s exact test, **\*\***p < 0.01, **\***p < 0.05.

Using these biomarker genes, we constructed a machine learning model to predict embryonic developmental competence (Fig. 6d). Support vector machine (SVM) was employed, and gene combinations and model parameters were systematically optimized. Model performance was evaluated using prediction accuracy, Area Under the Curve (AUC), and misclassification rate. The model using expression of *Zmym6*, *Sipa1*, and *Dppa3* achieved the best performance, with a prediction accuracy of 72%, an AUC of 0.70, and a cross-validation error below 0.05 (Fig. 6e, Extended Data Fig. 8a). We further validated the model in a prospective cohort. The developmental rates were 27% in the non-predicted group, 64% in the Good-predicted group, and 19% in the Bad-predicted group, with the Good group showing a significantly higher developmental rate compared to the others (Fig. 6f). The overall prediction accuracy reached 77.4% ± 10.5% (mean ± SD), with a sensitivity of 64.2% ± 22.8% and specificity of 80.8 ± 13.9%. Notably, the model demonstrated high specificity in identifying low-quality embryos, indicating its utility for excluding embryos with poor developmental potential. Furthermore, absolute quantification by digital PCR confirmed that *Sipa1* and *Zmym6* were consistently and reliably detected in both zygotes and PB2s, supporting their suitability as robust targets for minimally invasive evaluation (Extended Data Fig. 8b). The variable expression of those biomarkers across zygote and PB samples were also observed (Extended Data Fig. 8b, the range of 420 to 1,955 transcripts for *Sipa1* and 344 to 1,569 transcripts for *Zmym6* in zygotes). Finally, this prediction model was used to improve the development to term. The advantage of this system is that the developmental competence can be evaluated as early as the zygote stage: the outcome can be obtained within 12 hours after fertilization. Thus, 2-cell embryo transfer experiments using embryos classified by the prediction model were performed within 24 hours after IVF. These embryo transfer experiments demonstrated that only the Good-predicted group resulted in implantation, and live births were observed exclusively in the Good group, whereas no implantation or offspring were detected in the Bad-predicted group (Fig. 6g).

We used DBA/2 mice, known to be low developmental rate to term ^36^, for this embryo transfer experiments, in order to discern the effect of our prediction system. PB-biopsied embryos developed into pups at an efficiency of 7.6%, but this was significantly improved by our prediction system (Extended Data Fig. 8c). These findings indicate that our model can effectively exclude embryos with low developmental potential while enriching those with a high potential for successful development and live birth. Thus, we develop a maternal transcript-based rapid prediction method to reduce the risk of miscarriage.

## Discussion

Our study establishes a conceptual and technical framework for linking maternal transcript abundance to embryonic developmental potential by leveraging developmentally dispensable polar bodies. By demonstrating that the PB2 transcriptome closely mirrors that of its sibling zygote, we provide solid evidence that PBs can serve as a proxy for assessing maternally inherited molecular states without compromising embryo viability. Furthermore, we identify *Sipa1* and *Zmym6* as biomarkers to predict embryonic development as early as the zygote stage. Our prediction model enabled us to select zygotes that are likely to develop into blastocysts and even pups. Based on our findings, we propose that our biomarker transcript-based prediction can help reduce implantation failures and miscarriage rates, the major barriers to human infertility treatment. This finding extends the current use of PB to detect chromosomal aberrations in the corresponding oocyte ^37^.

Our work provides an experimental platform for identifying maternal transcripts associated with embryonic developmental competence. Although global transcriptomic profiles between embryos with high and low developmental potential were highly similar, a small set of DEGs was sufficient to discriminate between these groups. This observation is consistent with emerging evidence that early embryogenesis is governed by subtle quantitative differences in key regulatory factors ^38^. Among the candidate markers identified, *Zmym6* and *Sipa1* showed robust predictive performance across species. Notably, *Sipa1*, signal-induced proliferation-associated gene 1, is a transcription factor that regulates multiple genes involved in signal transduction, DNA synthesis, cell adhesion, and cell migration ^39^. In particular, crucial functions of *Sipa1* have been demonstrated in cancer metastasis ^40^. Maternal accumulation of *Sipa1* mRNA was observed in *Xenopus laevis*, with its RNA predominantly expressed in various sensory organs during later embryonic development ^41^. However, the roles of *Sipa1* in oogenesis and early embryonic development have never been investigated. *Zmym6*, zinc finger MYM-type containing 6, has been predicted to have DNA binding activity and might be involved in cytoskeleton organization. However, its roles have not been characterized, and further validation could reveal the molecular mechanisms underlying embryo quality. Thus, our experimental system enabled the identification of unprecedented biomarkers to assess embryonic quality.

Importantly, our findings also highlight a conceptual limitation of current embryo selection strategies. Morphological grading and even chromosomal screening approaches, such as PGT-A, do not fully capture the molecular state of embryos ^1,2^. In contrast, maternal transcript profiling provides a direct readout of the regulatory landscape that governs early development. Such developmental states may reflect coordinated transcriptional programs associated with early embryogenesis ^42^. Recent advances in time-lapse imaging and AI-based embryo scoring ^3^, as well as metabolomic profiling of culture media ^4^, underscore the growing interest in dynamic and non-invasive biomarkers. Our approach complements these methods by offering a molecular dimension that is both predictive and mechanistically informative.

From a broader perspective, our results support a model in which early embryogenesis can be understood as a tightly constrained dynamical system initiated by maternal inputs. Rather than being determined by discrete binary states, developmental competence may emerge from the trajectory of molecular states over time, shaped by the coordinated regulation of maternal transcripts, translational control, and chromatin remodeling ^43,44^. In this context, the maternal transcriptome captured in polar bodies may represent the initial conditions of this dynamical system. Small deviations in these initial conditions could propagate over time, ultimately determining developmental outcomes. This developmental strategy aligns with recent efforts to model biological processes as trajectory-based systems ^45,46^ and suggests that integrating transcriptomic, temporal, and morphological data may further enhance predictive accuracy.

Clinically, the application of this method to ART holds significant promise. Because polar body biopsy is already compatible with existing clinical workflows ^15^, incorporating transcriptomic analysis could enable rapid and minimally invasive assessment of developmental competence prior to transfer. Importantly, our prediction model achieved high accuracy in forecasting blastocyst development and even a high developmental rate to term. In human ART, our approach could reduce the transfer of embryos with low developmental potential, potentially leading to lower miscarriage rates and improved implantation success. Additionally, this strategy may be particularly valuable in cases where early embryo transfer using cleaving preimplantation embryos is preferred or when PGT-A results are inconclusive. For livestock propagation, our method enables assessment of developmental competence and demonstrates the feasibility of detecting selected expressed maternal trait-associated variants before embryo transfer, potentially enhancing breeding program efficiency.

Despite these advances, several limitations should be acknowledged. First, the number of transcripts detectable in polar bodies is biased toward highly abundant maternal RNAs, potentially limiting the detection of low-abundance but functionally critical regulators. Second, although our model performs well in predicting blastocyst formation, its ability to predict implantation and live birth outcomes remains to be validated in large-scale clinical studies. Third, the biological mechanisms linking specific maternal transcripts to developmental competence require further functional validation, including loss- and gain-of-function experiments. Lastly, cross-species comparison revealed only several genes with consistent expression patterns between mouse and bovine PB-outcome contrasts (Extended Data Fig. 4c), suggesting that a species-tuned biomarker panel may be required for each application context. Looking forward, several promising directions emerge from this work. The integration of maternal transcript profiling with existing non-invasive technologies, such as time-lapse imaging and spent media metabolomics, could yield multimodal prediction models with superior accuracy. Advances in microfluidic technologies and rapid RNA quantification methods, such as digital droplet PCR or isothermal amplification, may enable real-time PB transcript analysis within the time constraints of IVF cycles. Furthermore, as our understanding of maternal-effect genes and their regulatory networks deepens ^24^, the set of predictive transcripts can be refined to enhance specificity and sensitivity. Finally, the application of single-cell multi-omics approaches to PBs—integrating transcriptomic, epigenomic, and proteomic data from the same sample—could provide an even more comprehensive assessment of oocyte and embryo quality.

In conclusion, our study demonstrates that maternal transcript profiling in polar bodies provides a powerful and minimally invasive approach for predicting embryonic developmental potential. By bridging molecular, developmental, and clinical perspectives, this work lays the foundation for next-generation embryo selection strategies and offers new insights into the maternal control of early embryogenesis.

## Methods

### Animals

Mice (BALB/c, DBA/2, ICR, MCH (ICR)) strains at 8-13 weeks of age were purchased from CLEA Japan, Japan SLC, Kiwa Laboratory Animals, or the Jackson Laboratory Japan. Mice were maintained in light-controlled, air-conditioned rooms. BALB/c, DBA/2, and ICR mice were used for *in vitro* fertilization (IVF) as described below. This study was carried out in strict accordance with the recommendations in the Guidelines of Kindai University for the Care and Use of Laboratory Animals and the Guidelines for Animal Experiments of the Faculty of Agriculture at Kyushu University. All experiments were approved by the Committee on the Ethics of Animal Experiments of Kindai University (Permit Number: KABT-31-003) and by the Animal Care and Use Committee of Kyushu University (approval numbers: A24-337). Mice were sacrificed by cervical dislocation, and every effort was made to minimize suffering and reduce the number of animals used in this study.

The study using cows was approved by the National Livestock Breeding Center located in Nishigo, Fukushima, Japan. All animals received humane care according to guideline numbers 6, 22, and 105 of the Japanese Guidelines for Animal Care and Use.

### Mouse IVF

Female ICR, BALB/c, and DBA/2 mice were injected intraperitoneally with 7.5 IU of pregnant mare serum gonadotropin (PMSG; Asuka Animal Health, Japan) and superovulated 48 hours later with 7.5 IU of human chorionic gonadotropin (hCG; Asuka Pharmaceutical). The cumulus-oocyte complexes were harvested from the oviducts 15 to 16 hours after hCG injection in pre-equilibrated human tubal fluid (HTF) medium. Sperm were collected from the caudal epididymis of male ICR mice. Sperm suspensions were incubated in HTF medium for 1 to 1.5 hours and fertilized at 37°C under 5% CO_2_. The sperm suspension was added to the oocyte culture medium, and morphologically normal fertilized eggs were collected 2 hours after fertilization. The recovered oocytes were treated with 0.1% hyaluronidase (Merck) in HTF medium at 37°C until the oocytes dispersed. Fertilized embryos were then cultured in potassium simplex optimized medium KSOM (ARK Resource) at 37°C under 5% CO_2_.

### Mouse embryo culture

For the group culture of mouse embryos, KSOM was prepared as 35 µL drops under liquid paraffin, and up to 15 fertilized embryos were placed in each drop. The time of insemination was designated as 0 h, and embryo development was monitored at 24 h intervals until 120 h post-insemination (hpi).

For individual culture, KSOM was prepared as 10 µL drops under liquid paraffin. Each fertilized embryo, including the one subjected to polar body biopsy, was placed individually into a single KSOM drop. To trace embryos and their sibling polar bodies, each drop was labeled with an identification number corresponding to the polar body biopsied from that embryo. Embryo culture was initiated at 0 h post-insemination, and developmental progression was assessed every 24 h until 120 hpi. 120 hpi.

### Embryo transfer

Two-cell stage embryos derived from IVF were subjected to embryo transfer experiments. Prior to embryo transfer, fertilized embryos that had undergone polar body biopsy were classified based on the prediction outcome using their sibling polar bodies and labeled either as Good PB–associated embryos or Bad PB–associated embryos, as described in the result section. Recipient female MCH (ICR) mice were prepared by mating with vasectomized males, and the presence of a vaginal plug was designated as 0.5 d post-coitus (dpc). Anesthesia was induced with an anesthetic mixture (medetomidine, midazolam, and butorphanol), and two-cell stage embryos were surgically transferred into the oviducts of pseudopregnant recipient females using standard procedures. In each recipient, embryos associated with Good PB and Bad PB were transferred to contralateral oviducts within the same recipient for inter-maternal variability. After embryo transfer, anesthesia was antagonized with atipamezole (Antisedan), and recipient females were allowed to recover in a warming environment. Postoperative care was provided according to standard animal care guidelines, and recipient females were subsequently maintained under standard housing conditions. Cesarean section and uterine analysis of implantation sites were performed in all recipients at 19.5 dpc.

### Collection of oocytes/zygotes and polar bodies

#### Isolation of second polar bodies and sample processing in the mouse

Second polar bodies were isolated from IVF-derived zygotes during 7–9 hpi using a laser-assisted micromanipulation system (XYclone; Hamilton Thorne). We collected polar bodies only when PBs were detached from their accompanying zygotes to avoid forcibly tearing those cells apart. The criteria of choosing PB2s are the following: (1) PB2s are associated with zygotes with two pronuclei, (2) Generally, PB1s frequently exhibit a damaged form and an intact PB was chosen as a PB2, (3) When two intact PBs were found in the perivitelline space, we did not use such zygotes for the sampling, (4) malformed (ex: too small or too big) PB2s were not collected for the analyses. Following micromanipulation, the manipulated zygotes were cultured individually as described above. If zygotes were analyzed by RNA-seq, the zona was removed using acidified Tyrode’s solution (Merck, T1788). After the zona removal, embryos were washed with PBS containing 0.1% BSA. Similarly, the isolated PBs were placed in 0.1% w/v BSA/PBS. Each zygote or PB in 1 μl of 0.1% w/v BSA/PBS was transferred into an individual PCR tube containing 9.5 μL of reaction buffer from SMART-seq v4 Ultra Low Input RNA Kit (Takara Bio, Japan, Z4889N) for a final volume of 10.5 μl, and then incubated at room temperature for 5 min. After cell lysis in the buffer, 1 μL of the solution was removed and replaced with 1 μL of the diluted ERCC RNA Spike-In Mix (diluted for 350 pg of total RNA in the case of an oocyte/zygote and 20 pg of total RNA in the case of a PB according to the vendor’s instructions; Thermo Fisher Scientific, 4456740). For RT-qPCR analyses, ERCC spike-in was not added. Those lysed samples were snap-frozen in liquid nitrogen and kept at −80 °C until use.

#### Classification of Good PB and Bad PB in the mouse

Fertilized embryos after PB biopsy were washed and individually cultured in KSOM until 120 hpi (5 days). Based on developmental outcomes, embryos that reached the blastocyst stage were classified as high-quality embryos, whereas embryos that failed to reach the blastocyst stage were classified as low-quality embryos. Second polar bodies linked to blastocyst-stage embryos were designated as Good PB, and those linked to developmentally arrested embryos were designated as Bad PB for Figs. 2 and 6. To note, we did not use embryos arrested at the very early stage of development, such as the 2-cell and 4-cell stages, as these embryos might be arrested due to critical developmental errors, such as chromosomal abnormalities, which could hamper the identification of biomarkers predicting development to term.

#### Isolation of the second polar bodies and sample processing in the cow

Bovine oocyte samples were collected from Black Japanese breed cows at the National Livestock Breeding Center, Japan, and ovum pick-up (OPU) was performed on both immature and mature oocytes. After oocyte collection, immature oocytes were cultured for 22 hours for *in vitro* maturation (IVM) in IVM medium (TCM199 supplemented with 0.002 AU FSH + 5% NCS [newborn calf serum]) at 38.5°C under 5% CO_2_ and humidity saturation. For IVF, oocytes were collected by OPU from cows that had undergone follicle growth stimulation and superovulation treatments. The collected oocytes at the metaphase II stage were then fertilized in the medium (BO solution supplemented with 5 mM hypothaurine + 5U/mL heparin) with frozen sperm (3.0×10^6^ /ml) that had been washed and treated, and after 6 hours in medium, the oocytes were washed and transferred to culture medium (CR1aa medium supplemented with 5% NCS + 0. 25 mg/ml linoleic acid albumin) and incubated at 38.5°C under 5% CO_2_, and humidity saturation incubator (see National Livestock Breeding center manual: https://www.nlbc.go.jp/assets/pdf/19-2.pdf). Similarly, *in vitro* matured oocytes were used for IVF. For both samples, 7-9 hours after fertilization, the zona pellucida of fertilized embryos was removed with 0.25% Actinase E-DPBS, washed with 0.1% PVA-PBS (-), and the embryos and second polar bodies were separated by repeated pipetting. Some of the zygotes and PBs were used for RNA-seq analyses, following the sampling method described in the “Isolation of second polar bodies and sample processing in the mouse” section. The zygotes after the PB biopsy were subjected to individual embryo culture, in which the manipulated zygote was transferred to WOW dish and cultured under time-lapse observation (CCMMULTI; Astec, Fukuoka, Japan). Of the embryos that reached the blastocyst stage, those were graded according to the criteria of the IETS manual ^27^; good quality blastocyst stage embryos with code 1 (excellent, good) were selected as “Very good group”; moderately good quality blastocyst stage embryos with code 2 were selected as “Moderately good group”. In addition, embryos arrested before the blastocyst stage were considered “Bad PB”. These samples were used for RNA-seq analyses.

#### Isolation of first polar bodies and sample processing in humans

Collection of human oocytes and associated first polar bodies was performed as previously described ^32^. Briefly, following follicular aspiration and denudation, oocytes were morphologically classified into germinal vesicle (GV), metaphase I (MI), and MII stages. Oocytes that reached the MII stage by 16:00 on the day of retrieval were defined as normally matured oocytes and used for infertility treatment. In contrast, oocytes that failed to reach the MII stage by 16:00 were further cultured. A few hours after incubation, some oocytes were matured to MII, and these mature oocytes were sampled for RNA-seq analysis (Fig. 5, MII-1-3). Those that matured to the MII stage the following morning were defined as delayed-maturation oocytes (Fig. 5, dMII-1-3). For MII oocytes, the zona pellucida was perforated using a laser-assisted system (LYKOS Clinical IVF laser system and Clinical Laser Software Legacy 5.12, Hamilton Thorne), and the first polar body was biopsied using a micropipette. The oocyte and its corresponding first polar body were individually collected to preserve their pairing information. All collected oocytes and polar bodies were subjected to RNA-seq analysis as described before ^32^.

Human oocyte and corresponding first polar body samples were obtained as surplus oocytes not used for infertility treatment with written informed consent under a protocol approved by the Clinical Research Ethics Review Committee of Mie University Hospital (H2018_066), and the samples and data acquisition studies were registered in the University Hospital Medical Information Network Clinical Trials Registry in Japan (UMIN000034811). In addition, the study was originally registered with the Japan Society of Obstetrics and Gynecology for research involving human sperm, ova, and fertilized ova. The use and re-analysis of the previously obtained human transcriptome data ^32^ were approved by the Clinical Research Ethics Review Committee of Mie University Hospital under a new protocol entitled “Exploratory study of genetic biomarkers for evaluating human oocyte quality” (H2024-009). This study was conducted in accordance with the Declaration of Helsinki.

### RNA-seq library preparation

For all samples, RNA-seq library preparation was performed using the same protocol. cDNA synthesis and amplification were performed using the SMART-Seq® v4 Ultra® Low Input RNA Kit (Takara Bio, Japan, Z4889N) according to the manufacturer’s instructions with minor modifications. Reverse transcription was followed by PCR amplification, with the number of cycles adjusted according to sample type. Amplified cDNA was purified using AMPure XP beads (Beckman Coulter), assessed using a Bioanalyzer with a High Sensitivity DNA Kit (Agilent), and normalized prior to library preparation using the Nextera XT DNA Library Preparation Kit (Illumina). The resulting libraries were quality-checked using a Bioanalyzer and quantified using Qubit (Thermo Fisher Scientific).

### Sequencing and data processing

Next-generation sequencing was performed as previously described ^32^. Briefly, Paired-end sequencing (51 bp + 25 bp) or single-end sequencing (75 bp) was performed using an Illumina NextSeq 500 system. Raw FASTQ files generated by Illumina sequencing were filtered for low-quality reads by sliding window trimming (window size of 10 bases, quality value < 20), and low-quality bases were trimmed from the ends of reads (quality value < 20) using Trimmomatic. Reads shorter than 20 bases and unpaired reads were discarded. Adapter sequences as well as poly(A), poly(T), and poly(G) tails were removed using Trim Galore (https://www.bioinformatics.babraham.ac.uk/projects/trim_galore/) and Cutadapt. Filtered reads were mapped to species-specific reference genomes using STAR. Mouse RNA-seq reads were mapped to the mouse reference genome (mm10), human RNA-seq reads were mapped to the human reference genome (hg19), and bovine RNA-seq reads were mapped to the Bos taurus reference genome (ARS-UCD1.2). Gene-level read counts were obtained using featureCounts.

### RNA-seq data analysis

RNA-seq reads were visualized using Integrative Genome Viewer (IGV). Raw read counts for each gene were obtained from mapped reads using featureCounts and used for downstream analyses. Genes with low expression levels were filtered prior to analysis, and genes with raw counts ≥ 5 in at least one sample were considered as expressed genes and retained for subsequent analyses. To evaluate transcriptomic similarity among samples, Pearson’s correlation coefficients were calculated using normalized expression values of genes defined as expressed (raw counts ≥ 5). Correlation coefficients were visualized as scatter plots and/or correlation heatmaps. For comparisons between PBs and corresponding oocytes or zygotes, correlation analysis was performed using genes commonly detected in both samples. Fragments per kilobase of exon per million mapped reads (FPKM) values were also calculated from mapped reads by normalizing to total counts. The overlap of expressed genes was shown as Venn diagrams using Draw Venn Diagram (https://bioinformatics.psb.ugent.be/webtools/Venn/).

Differentially expressed genes (DEGs) were identified using DESeq2 based on raw read count data obtained by featureCounts by normalizing with ERCC RNA Spike-In counts (padj < 0.05 or *P* value < 0.05 and |log2FC|>2 or |log2FC|>1). Volcano plots were generated using log2 fold changes and −log10-transformed P values or adjusted P values derived from DESeq2 analysis. Genes with adjusted P values < 0.05 were considered significant in almost all comparisons, except for Extended Fig. 3c, in which subtle differences in gene expression between *in vitro*- and *in vivo*-derived zygotes were shown as P values < 0.05. An unsupervised hierarchical clustering of read count values was performed using hclust in TCC (unweighted pair-group method with arithmetic mean, UPGMA). Principal-component analysis (PCA) of the global gene expression profile was performed using scikit-learn and the plots were depicted using plotly. Gene ontology terms showing over-representation of genes that were up- or down-regulated were detected using DAVID tools (*P* < 0.05). Significant GO terms were identified using the modified Fisher Exact P-values according to the DAVID functional annotation tool. Furthermore, the DEG gene list was compared with the categorized gene lists reported in the previous paper ^25^. Each gene list was further analyzed using Ingenuity Pathway Analysis (IPA; Qiagen, Redwood City, USA, https://www.qiagenbioinformatics.com/products/ingenuity-pathway-analysis). Using IPA, enriched canonical pathways, upstream transcriptional regulators, gene regulatory networks, and diseases and biological functions in the identified DEGs were investigated. In IPA, *P* values were obtained using the statistical framework implemented in the software package. In addition, GREAT (http://great.stanford.edu/public/html/) was used to predict functions of DEGs. Expression levels of DEGs were displayed in heatmaps, with z-scores calculated from FPKM values using Heatmapper.

Previously published RNA-seq data of human oocytes (GSM5073832 for MII-1 [MII_S18]; GSM5073833 for MII-2 [MII_S27]; GSM5073834 for MII-3 [MII_S32]; GSM5073835 for dMII-1 [dMII_S08]; GSM5073836 for dMII-2 [dMII_S21]; GSM5073837 for dMII-3 [dMII_S25]) were used in Fig. 5 to compare with PB transcriptomes.

### RT-qPCR

The collected PB samples in the lysis buffer were directly used for reverse transcription using gene-specific primer (GSP) together with Oligo dT and SuperScript® III Reverse Transcriptase (ThermoFisher Scientific; 18080044). Gene-specific reverse primers for RT were selected from the qPCR primer list (Extended Data Table S2). cDNA samples were analyzed in the StepOne Real-Time PCR System (Applied Biosystems). For Extended data Fig 7b, c, reverse transcriptase reaction and cDNA amplification were performed according to the protocol of SMART-Seq v4 Ultra Low Input RNA Kit (Takara: Z4889N), SMART-Seq HT kit (Takara: 634437), or SMART-Seq mRNA LP kit (Takara: Z4768N). Each sample was diluted 50-fold and used as a template for qPCR. We calculated the expression level of Good PB relative to Bad PB by dividing the qPCR quantity for Good PB by the average Bad PB quantity, setting the average Bad PB expression level to 1. We performed F- and T-tests to identify genes with differential expression between Bad PB and Good PB (Fig. 6c, Extended data Fig 7b-e).

### Digital PCR

The cDNA preparation for a single PB2 or zygote sample was performed as described above for the RT-qPCR step. Then the copy number (cp) of cDNA was measured by QuantStudio Absolute Q (Applied Biosystems). Ten ul rection mixtures were prepared for each sample according to the vendor’s recommendation. The cp values in the raw samples (per embryo and per polar body) were then calculated based on the volume used.

### Machine learning analysis

Machine learning analysis was conducted in two sequential steps.. First, combinations of expression data of candidate genes were systematically evaluated to identify an optimal feature set for discriminating Good PB and Bad PB. Second, a final prediction model was constructed using the selected feature set and applied to unlabeled polar body samples.

#### Feature set evaluation and selection

Prior to the construction of the final prediction model, combinations of candidate genes using their expression data in PBs were evaluated to identify an optimal feature set for discriminating Good PB and Bad PB. Four genes, namely *Cdk2ap1*, *Sipa1*, *Zmym6*, and *Dppa3*, quantified by single PB RT-qPCR were considered, and all possible combinations (15 subsets) were assessed. Feature subsets were evaluated using cross-validation to minimize overfitting and improve prediction performance across developmental outcomes ^47^. For each gene combination, a support vector machine (SVM) was trained and evaluated using cross-validation within the training dataset only. Model performance was assessed using multiple metrics, including accuracy, area under the receiver operating characteristic curve (AUC), sensitivity, specificity, and cross-validation loss. Test data were not used during feature selection. Based on these metrics, the most suitable gene combination was selected and used to train the final prediction model. The detailed comparison of all gene combinations and the selected optimal feature set are presented in the Results section. For each gene combination, model training and performance evaluation were carried out using identical sample sets, with differences only in the selected feature subsets. Performance comparison among gene combinations was conducted exclusively using cross-validation within the training dataset.

#### Model training with class imbalance correction

The final prediction model was constructed using SVM with a radial basis function (RBF) kernel implemented in MATLAB. To address class imbalance between Good PB and Bad PB samples, the Synthetic Minority Over-sampling Technique (SMOTE) ^48^ was applied exclusively to the training data prior to model fitting. Synthetic samples were generated using a k-nearest neighbors algorithm (k = 5) to balance class distributions. After resampling, feature values were standardized and used to train the SVM classifier. The SVM hyperparameters were fixed as follows: BoxConstraint = 10.83 and KernelScale = 0.45. To account for asymmetric misclassification risks, a cost matrix was introduced (Good PB misclassified as Bad PB = 4; Bad PB misclassified as Good PB = 6).

#### Classification and decision threshold

After model training, class scores were obtained for each sample using the trained SVM model. A decision threshold of 0.2 was applied to the Good PB class score to convert continuous scores into binary class labels. Samples with scores exceeding the threshold were classified as Good PB, whereas those below the threshold were classified as Bad PB.

#### Prediction of unlabeled PB samples

The trained SVM model was applied to unlabeled polar body datasets to predict whether each polar body was Good PB or Bad PB. Predicted class labels were assigned to each sample and used for downstream analyses, including evaluation of embryo developmental outcomes and embryo transfer experiments.

### Quantification and Statistical Analysis

Data on embryonic development were analyzed by the Chi-square test (Fig. 6f and Extended Data Fig. 2c, 2d, 8c) or Fisher’s exact test (Fig. 6g and Extended Data Fig. 8c); an appropriate test was decided depending on the expected values of the samples. Gene expression levels were compared by F- and T-test (Fig. 2c, 5c, 6c and Extended Data Fig. 1c, 1f, 7c, 7e), or ANOVA (Fig. 4d). All of the statistical details, including the number of embryos used for analyses and experimental or biological repeats, were described in figures and/or figure legends. A value of P < 0.05 was considered significant.

## DATA AVAILABILITY

The accession number for the RNA-seq data generated in this study and reported in this paper is GEO: GSE334135. Requests for additional program codes and data generated and/or analyzed during the current study should be directed to and will be fulfilled upon reasonable request by the corresponding author.

## AUTHOR CONTRIBUTIONS

K.Mi. conceived the project. K.Mi and A.I. designed the experiments. A.I., M.Y., Y.W., K.H., H.S., and K.Mi. performed mouse experiments. T.Y., H.M., H.Y., G.R., and K.Mi. performed bovine experiments. H.T and M.N. performed human experiments. A.K., H.K., and T.K. performed NGS analyses. K.Mi., A.I., and Y.W. wrote the manuscript. A.I., N.C., and K.Mi. performed bioinformatics analyses. K.Mi. edited the manuscript. K.Ma. provided the experimental tools. K.Mi supervised research.

## ACKNOWLEDGEMENTS

We thank Mr Tomomasa Tsukaguchi, Ms Manami Fukui, Ms Madoka Karatsu, Ms Ayumi Doi, Ms Fumi Arai and Ms Ayaka Nishira for technical support. We thank Mr Shinnosuke Tamura, Mr Yunosuke Yamamoto and Mr Shin Hongo for technical assistance in the bovine experiments, as well as the members of Animal Reproduction Technology Team at the National Livestock Breeding Center for animal management. We are also grateful to Dr Nobuhiko Yamauchi and Dr Kazuo Yamagata for their constructive advice. This research was supported by Japan Society for the Promotion of Science KAKENHI grant (grant numbers: JP19H05751, JP20K21376, JP23K27092, and JP25H02569 [to K. Mi.], by JST FOREST Program (grant no.: JPMJFR233F; to K. Mi), by JST CREST Program (grant no.:JPMJCR2572; to K. Mi), by Takeda Science Foundation (to K. Mi.), by The Naito Foundation grant (to K. Mi.), by AMED under Grant Number JP25yf0126004, and the Center for Clinical and Translational Research of Kyushu University, and by Young Researcher Support Project of Mie University to H.T. This work was supported by the Cooperative Research Grant of the Genome Research for BioResource, NODAI Genome Research Center, Tokyo University of Agriculture.

---

**Extended Data Fig. 1.**
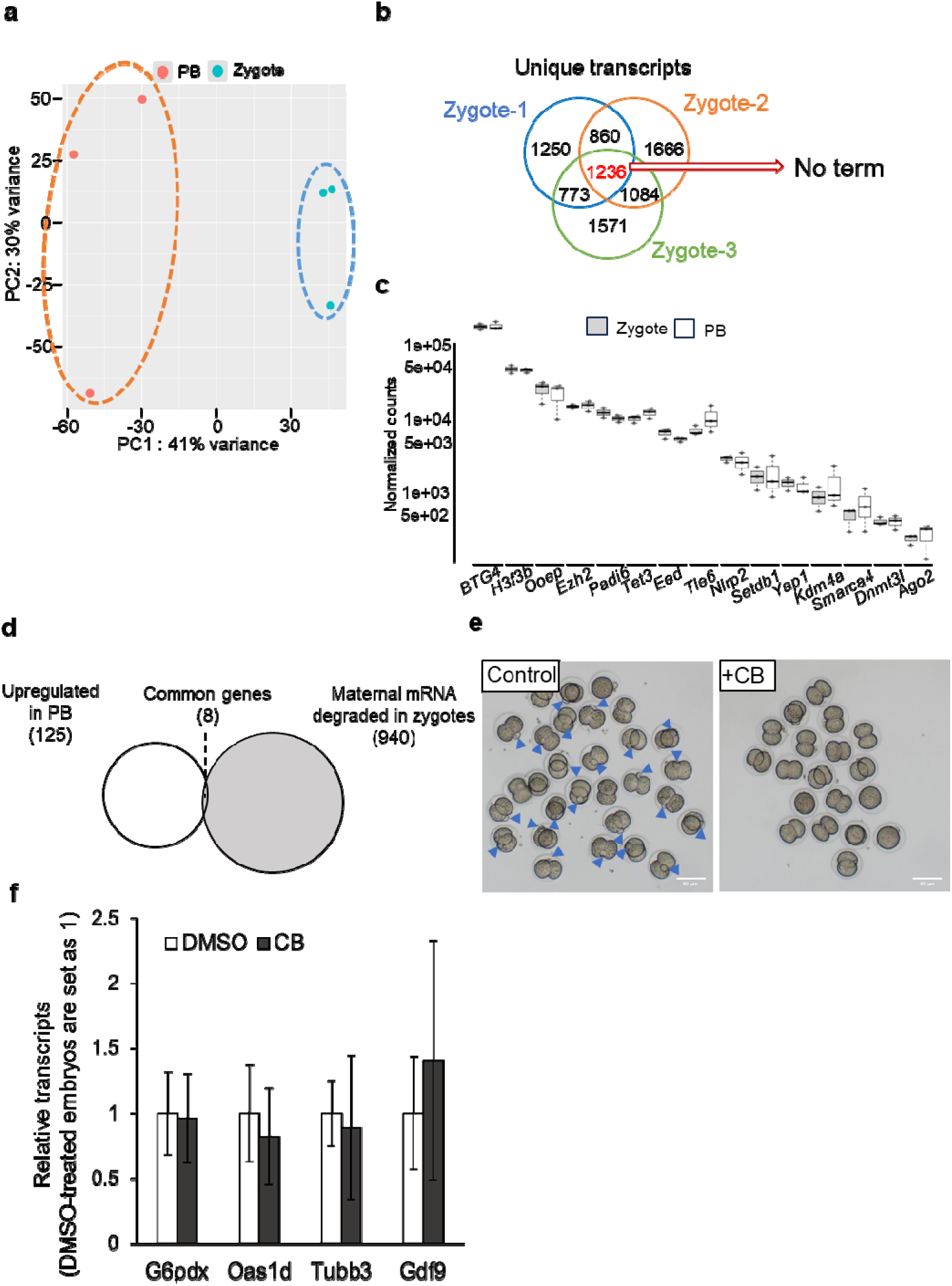
Characterization of transcriptional profiles of polar bodies in mouse. **a**, PCA of zygote and PB transcriptomes. **b**, A Venn diagram shows the overlap of unique transcripts in Fig. 1a, and no GO terms were found using the shared transcripts (1,236 transcripts) by GREAT analysis. **c**, Normalized transcript counts of maternal effect genes in mouse zygotes and PBs. The expression levels did not differ between zygotes and PBs, as determined by the T-test (P >0.05). **d**, A Venn diagram showing the overlap between upregulated transcripts in PB (Fig. 1f) and the maternal mRNAs that are degraded in zygotes ^25^. **e**, Cytochalasin B (CB)-treated 2-cell embryos and the control group (treated by DMSO). Release of PBs (blue arrowheads) was inhibited in the CB-treated group. **f**, RT-qPCR analyses of Cytochalasin B (CB) treated 2-cell embryo and the control group (treated by DMSO). Mean ± SD. P > 0.05 (T-test), N=3 biological replicates.

**Extended Data Fig. 2.**
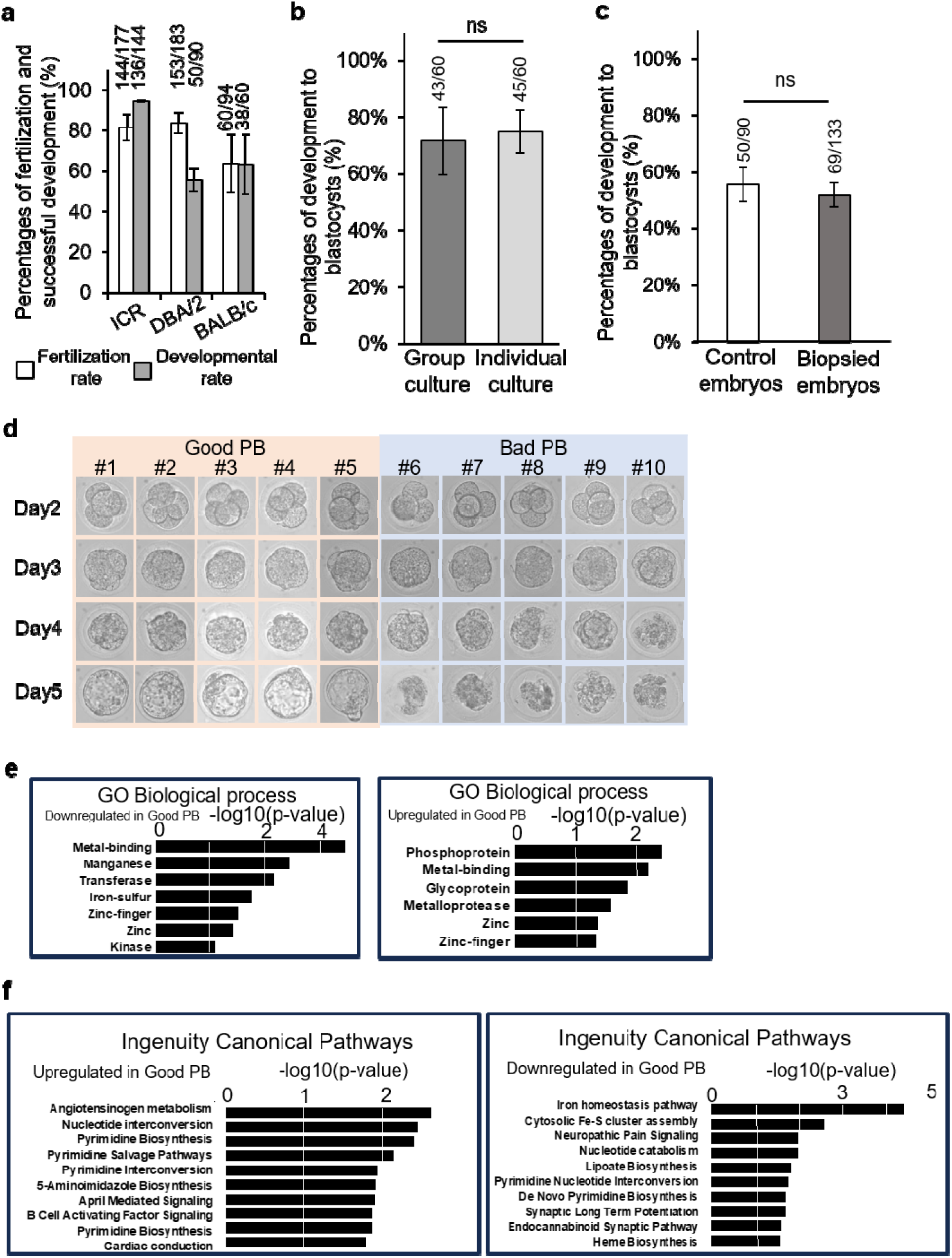
Establishment of a minimally invasive system to measure maternal transcripts, associated with the developmental potential, using PB2s. **a**, Fertilization and developmental rates to the blastocyst stage of the three mouse strains: ICR, DBA/2, and BALB/c. Mean ± SE. N = 3 experiments. The total number of embryos in each group is shown above. **b**, Developmental rates of DBA/2 mouse embryos cultured in a group way (one culture medium drop with dozens of embryos) or an individual way (one culture medium drop with one embryo). Mean ± SE. N = 3 experiments. The total number of embryos in each group is shown above. ns, as judged by the Chi-square test. **c**, Developmental rates of DBA/2 mouse embryos with or without PB2 biopsy. Mean ± SE. N = 3 experiments. The total number of embryos in each group is shown above. ns, as judged by the Chi-square test. **d**, Bright-field images of preimplantation development of embryos that were grouped into Good PB and Bad PB in Fig. 2. **e**, GO biological process enrichment analyses of upregulated genes in Good- and Bad-PB samples. **f**, Ingenuity Pathway Analysis (IPA) of upregulated genes in good- and Bad-PB samples.

**Extended Data Fig. 3.**
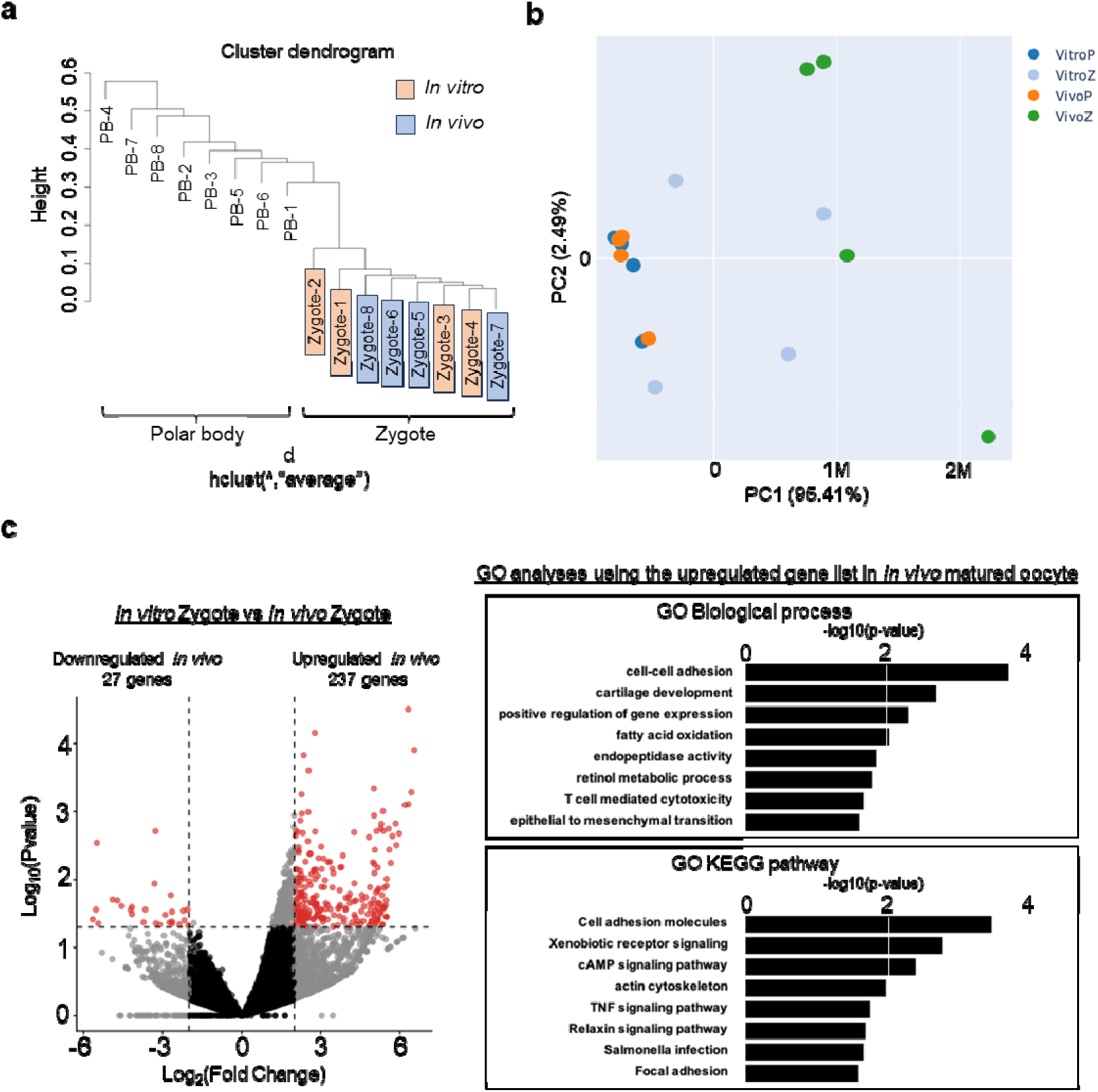
Characterization of transcriptomes of bovine zygotes and PB2s derived from *in vitro* matured and *in vivo* matured oocytes. **a**, Cluster dendrogram of transcriptomic profiles from zygotes and their sibling PB2s derived from different maturation conditions (*in vitro* and *in vivo* matured). Samples cluster primarily by cell type (PB vs zygote). Color coding indicates sample groups (*in vitro* and *in vivo* matured). **b**, PCA of transcriptomes from zygotes and PB2s under *in vitro* and *in vivo* maturation conditions. **c**, A volcano plot showing DEGs between zygotes derived from *in vitro* and *in vivo* maturation (|FC|≧2, P value<0.05). Dashed lines indicate significant difference and fold-change thresholds. The right graph summarizes the results of GO analyses using the upregulated gene list in the *in vivo* zygote group (237 genes).

**Extended Data Fig. 4.**
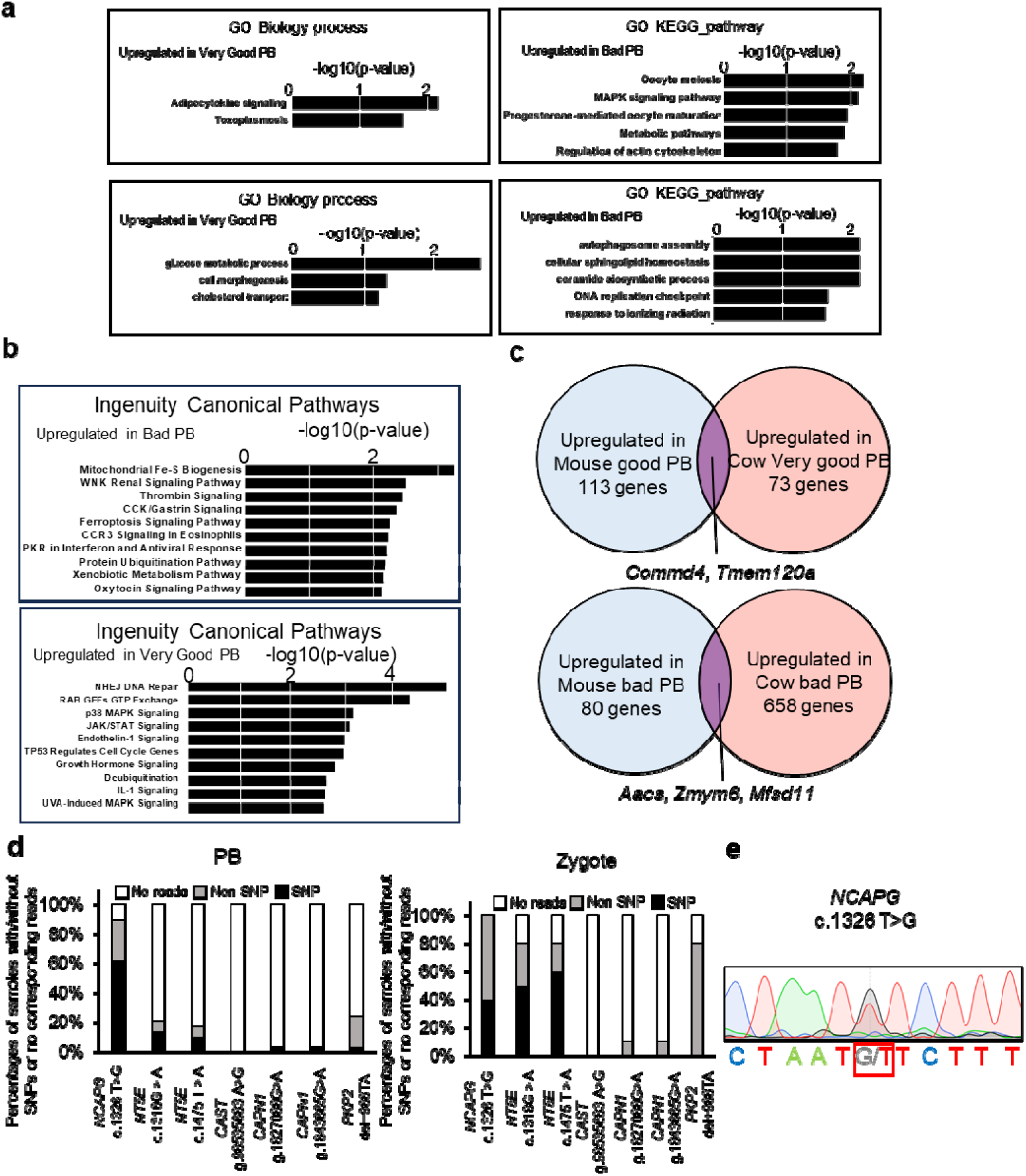
Characterization of our minimally invasive method to measure biomarker traits in bovine PBs. **a**, GO biological process and KEGG pathway enrichment analyses of genes upregulated in very Good/Bad PB samples. **b**, IPA of genes upregulated in very Good/Bad PB samples. **c**, Venn diagrams showing the overlap of DEGs identified in mouse and bovine PB analyses (Log_2_FC>2, Padj<0.05 or Log_2_FC<-2, Padj<0.05). **d**, Detection rates of SNPs in bovine PB and zygote RNA-seq reads, related to carcass weight (*NCAPG*) ^28^, Inosinic acid (*NT5E*) ^49^, meat tenderness (*CAPN1* and *CAST*) ^29,30^, and conception rate (*PKP2*) ^50^. **e**, Sequencing analyses of the bovine *NCAPG* gene using genomic DNA obtained from a calf (Fig. 4h).

**Extended Data Fig. 5.**
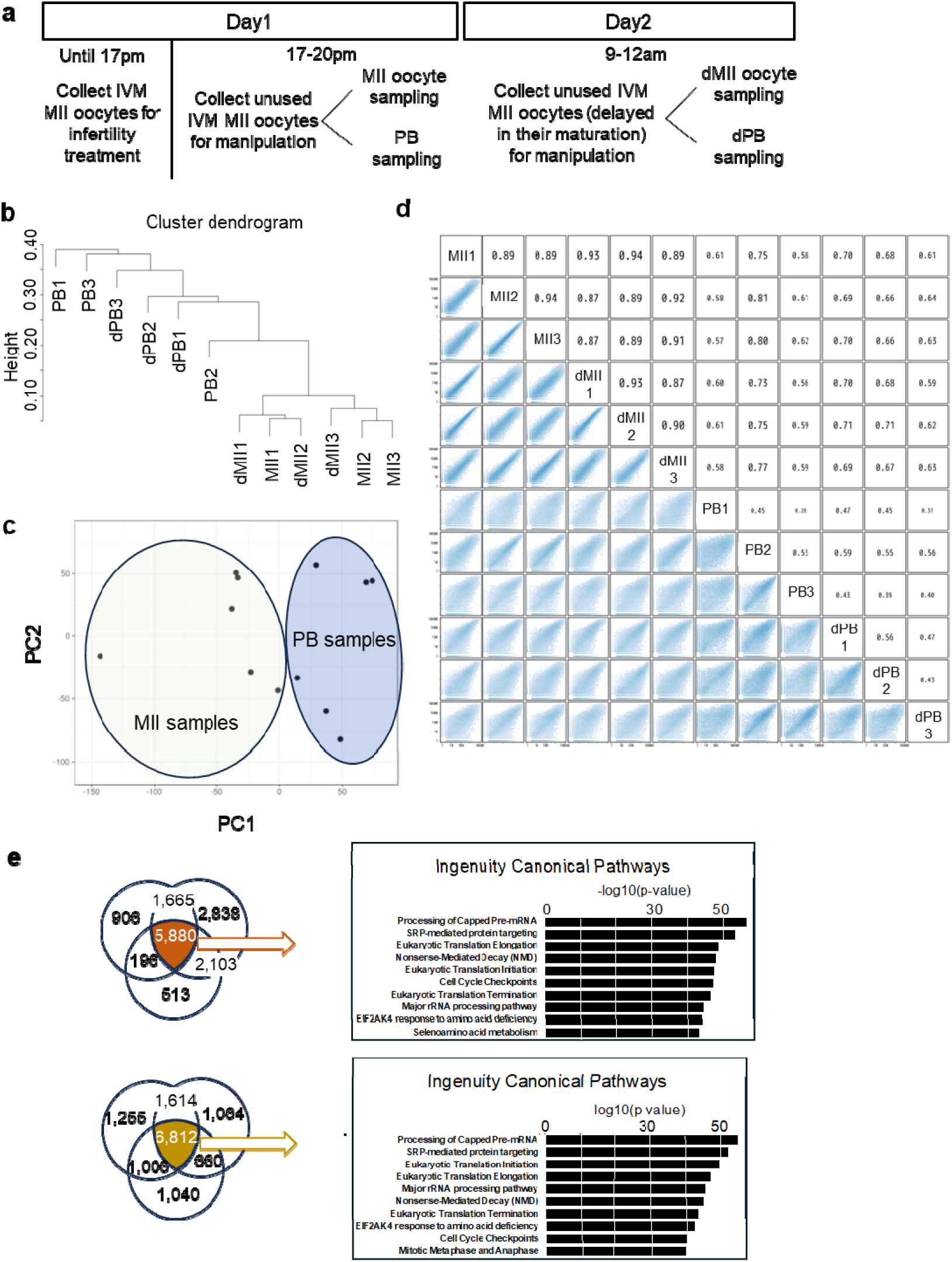
Transcriptomic relationships between human MII oocytes and polar bodies. **a**, Workflow for human MII oocyte / PB sampling. **b**, A clustering dendrogram showing relationships among MII oocytes, delayed MII (dMII) oocytes, and corresponding polar bodies. **c**, PCA of transcriptomes from PBs and MII oocytes. **d**, Correlation matrix of transcript expression levels across all samples. The upper triangle shows Spearman correlation coefficients, and the lower triangle shows scatter plots of gene expression comparisons. **e**, IPA of the shared transcripts (same as Fig. 5a), highlighting the common canonical pathways.

**Extended Data Fig. 6.**
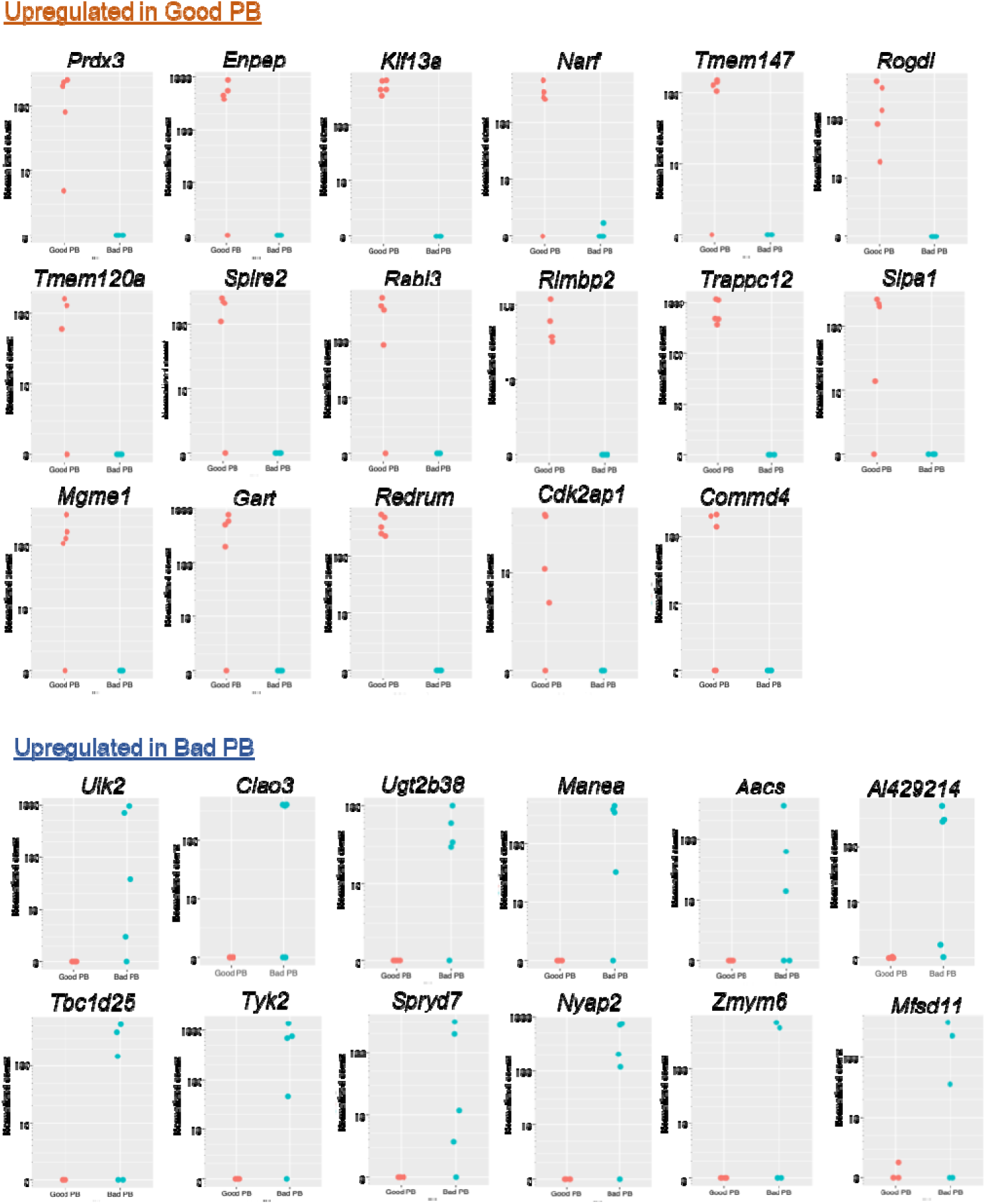
Expression of candidate biomarker transcripts in mouse Good and Bad PBs. Normalized count values were compared between Good and Bad PBs, as determined by RNA-seq analysis.

**Extended Data Fig. 7.**
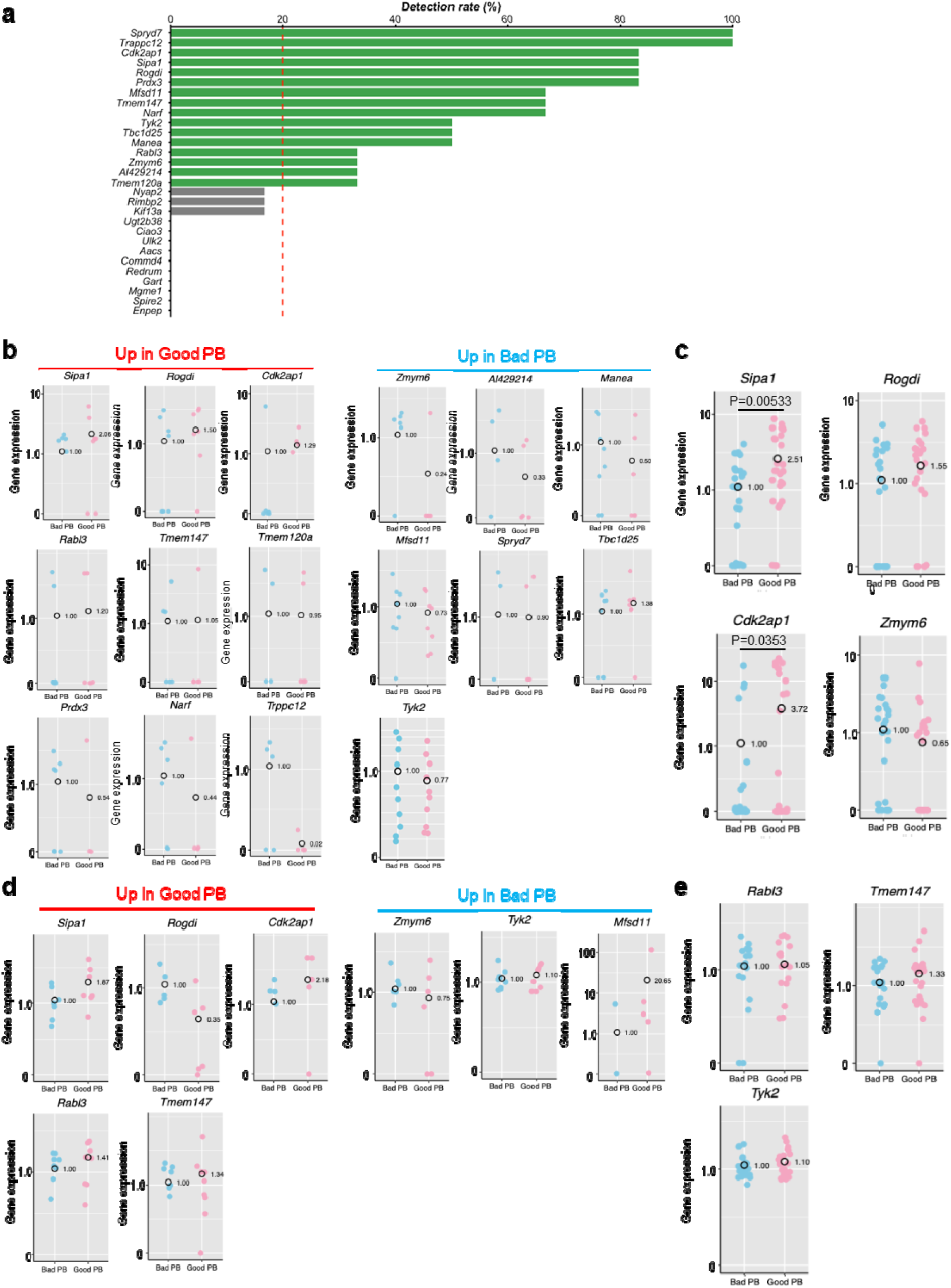
Screening of biomarker transcripts in mouse PB2s. **a**, Percentages of successful detection of each gene by single PB RT-qPCR. We set the threshold at 20% of the detection rate, and genes whose transcripts were not detected in more than 80% of PB samples were not further validated. **b,c**, RT-qPCR was performed using the cDNA-amplified Good and Bad PB samples. The means of Bad PBs were set as 1. Each spot corresponds to each PB sample. T-test was performed, and when significant differences were found, P values are indicated. **d,e**, Relative expression levels of candidate biomarkers, as judged by single PB RT-qPCR. Each spot corresponds to each PB sample. P >0.05 (T-test).

**Extended Data Fig. 8.**
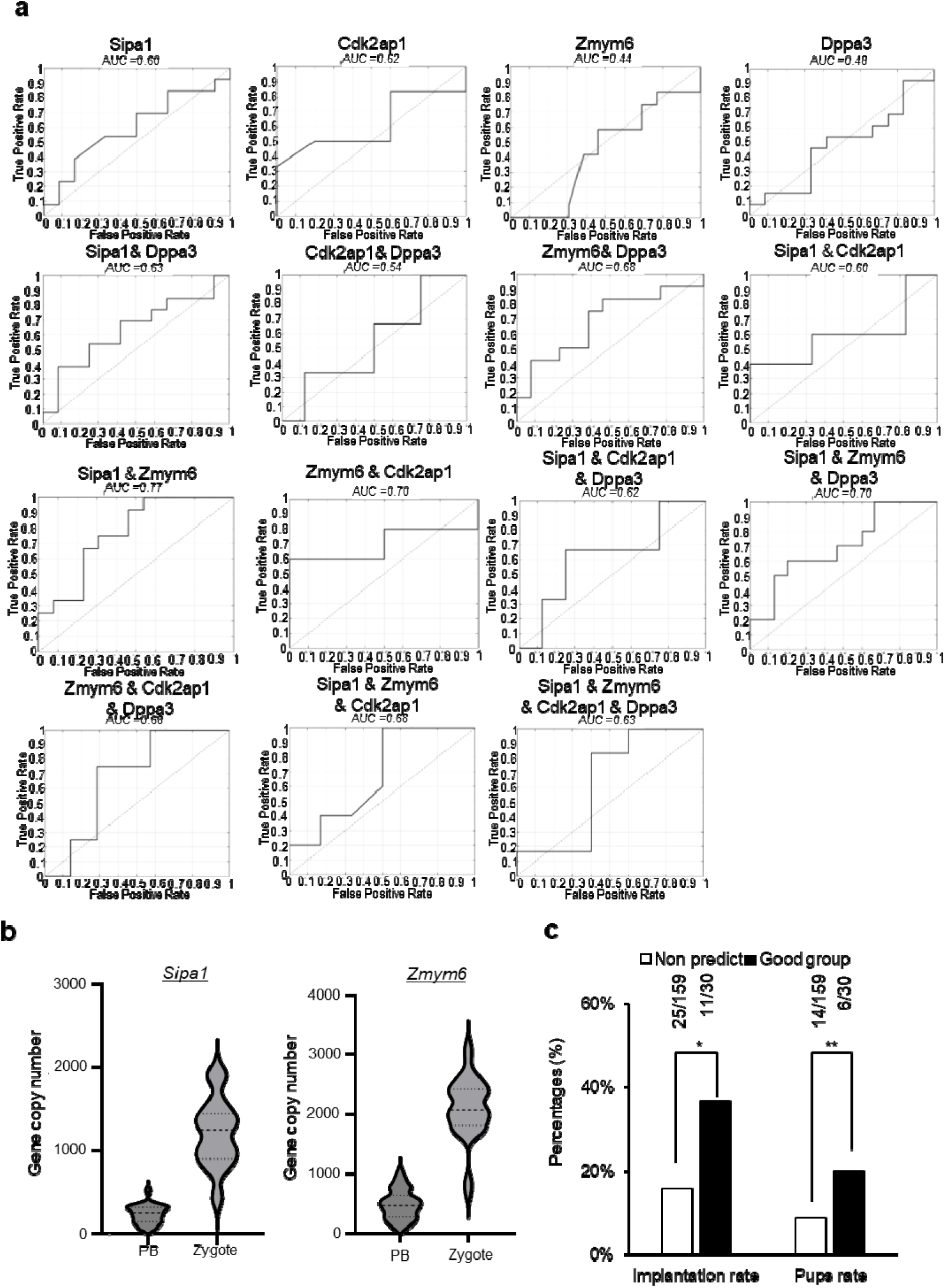
Evaluation of predictive performance across individual gene combinations. **a**, Receiver operating characteristic (ROC) curves for classification of developmental competence of embryos using individual biomarker candidate genes or gene combinations. Each panel represents an independent model, with area under the curve (AUC) values indicated. **b**, Violin plots showing gene copy number of *Sipa1* and *Zmym6* in single PB and zygote, as determined by digital PCR. N = 28 embryos for *Sipa1* and *Zmym6*. **c,** The effect of PB-based developmental prediction on the implantation rate and litter size. *Fisher’s exact test, P< s0.05; **Chi-square test, P< 0.01.

## REFERENCES

1. Munné, S., Kaplan, B., Frattarelli, J. L., & others. Preimplantation genetic testing for aneuploidy versus morphology as selection criteria for single frozen-thawed embryo transfer in good-prognosis patients: a multicenter randomized clinical trial. Fertil Steril 112, 1071–1079.e7 (2019).

2. Greco, E., Minasi, M. G. & Fiorentino, F. Healthy Babies after Intrauterine Transfer of Mosaic Aneuploid Blastocysts. N Engl J Med 373, 2089–2090 (2015).

3. Bori, L., Meseguer, F., Valera, M. A., & others. The higher the score, the better the clinical outcome: retrospective evaluation of automatic embryo grading as a support tool for embryo selection in IVF laboratories. Hum Reprod 37, 1148–1160 (2022).

4. Alizadeh Moghadam Masouleh, A., Eftekhari-Yazdi, P., & others. Embryo metabolism as a novel non-invasive preimplantation test: nutrients turn over and metabolomic analysis of human spent embryo culture media (SECM). Hum Reprod Update 31, 405–444 (2025).

5. Tadros, W. & Lipshitz, H. D. The maternal-to-zygotic transition: a play in two acts. Development 136, 3033–3042 (2009).

6. Yan, L., Yang, M., Guo, H., & others. Single-cell RNA-Seq profiling of human preimplantation embryos and embryonic stem cells. Nat Struct Mol Biol 20, 1131–1139 (2013).

7. Li, L., Zheng, P. & Dean, J. Maternal control of early mouse development. Development 137, 859–870 (2010).

8. Fan, Y. et al. Aberrant expression of maternal Plk1 and Dctn3 results in the developmental failure of human in-vivo- and in-vitro-matured oocytes. Sci Rep 5, 8192 (2015).

9. Tong, Z. B., Gold, L., Pfeifer, K. E., & others. Mater, a maternal effect gene required for early embryonic development in mice. Nat Genet 26, 267–268 (2000).

10. Xu, Y., Shi, Y., Fu, J., & others. Mutations in PADI6 Cause Female Infertility Characterized by Early Embryonic Arrest. Am J Hum Genet 99, 744–752 (2016).

11. Bebbere, D., Masala, L., Albertini, D. F. & Ledda, S. The subcortical maternal complex: multiple functions for one biological structure? J Assist Reprod Genet 33, 1431–1438 (2016).

12. Zhao, C. et al. A comprehensive human embryo reference tool using single-cell RNA-sequencing data. Nat Methods 22, 193–206 (2025).

13. Proks, M., Salehin, N. & Brickman, J. M. Deep learning-based models for preimplantation mouse and human embryos based on single-cell RNA sequencing. Nat Methods 22, 207–216 (2025).

14. Hou, Y. et al. Genome analyses of single human oocytes. Cell 155, 1492–1506 (2013).

15. Verpoest, W., Staessen, C., Bossuyt, P. M., & others. Preimplantation genetic testing for aneuploidy by microarray analysis of polar bodies in advanced maternal age: a randomized clinical trial. Hum Reprod 33, 1767–1776 (2018).

16. Ma, H. et al. Functional Human Oocytes Generated by Transfer of Polar Body Genomes. Cell Stem Cell 20, 112–119 (2017).

17. Wang, T. et al. Polar body genome transfer for preventing the transmission of inherited mitochondrial diseases. Cell 157, 1591–1604 (2014).

18. Reich, A., Klatsky, P., Carson, S. & Wessel, G. The Transcriptome of a Human Polar Body Accurately Reflects Its Sibling Oocyte. J Biol Chem 286, 40743–40749 (2011).

19. Jiao, Z.-X. & Woodruff, T. K. Detection and quantification of maternal-effect gene transcripts in mouse second polar bodies: potential markers of embryo developmental competence. Fertil Steril 99, 2055–2061 (2013).

20. Jin, H. et al. The second polar body contributes to the fate asymmetry in the mouse embryo. Natl Sci Rev 9, nwac003 (2022).

21. Qing, T., Yu, Y., Du, T. & Shi, L. mRNA enrichment protocols determine the quantification characteristics of external RNA spike-in controls in RNA-Seq studies. Sci. China Life Sci. 56, 134–142 (2013).

22. Jiang, L. et al. Synthetic spike-in standards for RNA-seq experiments. Genome Res 21, 1543–1551 (2011).

23. Hu, Y. et al. Nascent transcriptome of embryonic genome activation reveals a regulatory axis linking transcriptional priming to early lineage specification in mouse embryos. Nucleic Acids Res 54, gkag173 (2026).

24. Mitchell, L. E. Maternal effect genes: Update and review of evidence for a link with birth defects. HGG Adv 3, 100067 (2021).

25. Park, S.-J., Shirahige, K., Ohsugi, M. & Nakai, K. DBTMEE: a database of transcriptome in mouse early embryos. Nucleic Acids Res 43, D771–776 (2015).

26. Yanez, L. Z., Han, J., Behr, B. B., Pera, R. A. R. & Camarillo, D. B. Human oocyte developmental potential is predicted by mechanical properties within hours after fertilization. Nat Commun 7, 10809 (2016).

27. Manual of the International Embryo Transfer Society: A Procedural Guide and General Information for the Use of Embryo Transfer Technology, Emphasizing Sanitary Procedures. (The Society, Savory, Ill, 1998).

28. Setoguchi, K., et al. The SNP c.1326T>G in the non-SMC condensin I complex, subunit G (NCAPG) gene encoding a p.Ile442Met variant is associated with an increase in body frame size at puberty in cattle. Anim Genet 42, 650–655 (2011).

29. Page, B. T. et al. Evaluation of single-nucleotide polymorphisms in CAPN1 for association with meat tenderness in cattle. J Anim Sci 80, 3077–3085 (2002).

30. Calvo, J. H. et al. A new single nucleotide polymorphism in the calpastatin (CAST) gene associated with beef tenderness. Meat Sci 96, 775–782 (2014).

31. Shirasawa, H. & Terada, Y. In vitro maturation of human immature oocytes for fertility preservation and research material. Reprod Med Biol 16, 258–267 (2017).

32. Takeuchi, H. et al. Single-cell profiling of transcriptomic changes during in vitro maturation of human oocytes. Reprod Med Biol 21, e12464 (2022).

33. Feng, R., Sang, Q., Kuang, Y., & others. Mutations in TUBB8 and Human Oocyte Meiotic Arrest. N Engl J Med 374, 223–232 (2016).

34. Pfender, S., Kuznetsov, V., Pleiser, S., Kerkhoff, E. & Schuh, M. Spire-type actin nucleators cooperate with Formin-2 to drive asymmetric oocyte division. Curr Biol 21, 955–960 (2011).

35. Modzelewski, A. J. et al. A mouse-specific retrotransposon drives a conserved Cdk2ap1 isoform essential for development. Cell 184, 5541–5558.e22 (2021).

36. Nishizono, H., Uno, K. & Abe, H. Cleavage Speed and Blastomere Number in DBA/2J Compared with C57BL/6J Mouse Embryos. J Am Assoc Lab Anim Sci 56, 11–17 (2017).

37. Oberle, A. & Feichtinger, M. Polar body-based PGT-A: not dead yet? A step forward back to the roots of PGT-A. Reproductive BioMedicine Online 50, 104430 (2025).

38. Wu, D. & Dean, J. Maternal factors regulating preimplantation development in mice. Curr Top Dev Biol 140, 317–340 (2020).

39. Guo, L. et al. A novel transcription factor SIPA1: identification and verification in triple-negative breast cancer. Oncogene 42, 2641–2654 (2023).

40. Park, Y.-G. et al. Sipa1 is a candidate for underlying the metastasis efficiency modifier locus Mtes1. Nat Genet 37, 1055–1062 (2005).

41. Rothe, M., Monteiro, F., Dietmann, P. & Kühl, S. J. Comparative expression study of sipa family members during early Xenopus laevis development. Dev Genes Evol 226, 369–382 (2016).

42. Wang, T. et al. Single-cell multi-omics profiling of human preimplantation embryos identifies cytoskeletal defects during embryonic arrest. Nat Cell Biol 26, 263–277 (2024).

43. Wyatt, C. D. R. et al. A developmentally programmed splicing failure contributes to DNA damage response attenuation during mammalian zygotic genome activation. Sci. Adv. 8, eabn4935 (2022).

44. Takada, Y. et al. Mature mRNA processing that deletes 3′ end sequences directs translational activation and embryonic development. Sci. Adv. 9, eadg6532 (2023).

45. Qu, R. et al. Gene trajectory inference for single-cell data by optimal transport metrics. Nat Biotechnol 43, 258–268 (2025).

46. Li, G. et al. Sceptic: pseudotime analysis for time-series single-cell sequencing and imaging data. Genome Biol 26, 209 (2025).

47. Pedregosa, F. et al. Scikit-learn: Machine Learning in Python. J. Mach. Learn. Res. 12, 2825–2830 (2011).

48. Chawla, N. V., Bowyer, K. W., Hall, L. O. & Kegelmeyer, W. P. SMOTE: Synthetic Minority Over-sampling Technique. Journal of Artificial Intelligence Research 16, 321–357 (2002).

49. Uemoto, Y. et al. Effect of two non-synonymous ecto-5’-nucleotidase variants on the genetic architecture of inosine 5’-monophosphate (IMP) and its degradation products in Japanese Black beef. BMC Genomics 18, 874 (2017).

50. Sugimoto, M. et al. Genetic variants related to gap junctions and hormone secretion influence conception rates in cows. Proceedings of the National Academy of Sciences 110, 19495–19500 (2013).

